# Ongoing hybridisation among clownfishes: the genomic architecture of the Kimbe Bay hybrid zone

**DOI:** 10.1101/2024.03.10.584293

**Authors:** Sarah Schmid, Diego A. Hartasánchez, Ashton Gainsford, Geoffrey P. Jones, Nicolas Salamin

## Abstract

Hybrid zones – locations where genetically distinct lineages interact and reproduce – are remarkable resources for exploring the evolutionary trajectory of species. Not only can we learn from hybrid zones about the mechanisms of speciation and how reproductive isolation is maintained, but we can also study their impact on evolutionary processes. Thanks to the advent of next-generation sequencing, we are now able to gain new insight into the structure of hybrid genomes and the factors influencing the outcome of hybridisation. Here, we focus on the Kimbe Bay hybrid zone, a narrow region in the Pacific Ocean where two species of clownfish – *Amphiprion chrysopterus* and *A. sandaracinos* – hybridise and give rise to the hybrid *A. leucokranos*. Based on whole-genome sequencing, we found that the hybrid zone is mainly composed of first-generation hybrids, the first evidence of F2 hybrids in the wild and early backcrosses with *A. sandaracinos*. The recurrent backcrossing with one of the parental species might lead to adaptive introgression, with few adaptive introgressed loci from *A. chrysopterus* integrated into the *A. sandaracinos* genomic background. This study builds upon the growing literature body relative to the evolutionary outcomes of hybridisation and its importance in the evolution of many species.

## INTRODUCTION

More than 30 years ago, the potential of hybrid zones to investigate the evolutionary history of species was already recognised in Hewitt’s and Harrison’s seminal works, who qualified them as “natural laboratories for evolutionary studies” (Hewitt 1988) or “windows on evolutionary process” (Harrison 1990). Today, the numerous hybrid zones studied have proved them right and have remarkably contributed to a remarkably improved understanding of species’ origin, maintenance and collapse (notably reviewed in Feder et al. 2012; Shurtliff 2013; Abbott et al. 2013; Abbott 2017)

Hybrid zones consist of geographic areas where divergent lineages exchange genetic variants through the process of hybridisation (Barton and Hewitt 1985; Harrison and Harrison 1993; Gompert and Buerkle 2016). They are an ideal focal point to study the interaction between divergent genomic backgrounds, which might help elucidate the mechanisms underlying speciation and eventually lead to the identification of candidate genes contributing to reproductive isolation (Hewitt 2001; Harrison and Larson 2014). Hybrid zones also experience ongoing evolutionary processes (Arnold 1997; Abbott et al. 2013); they can act as conduits for adaptive alleles and enable them to cross species boundaries (Pardo-Diaz et al. 2012), or increase reproductive isolation by reinforcement or coupling of incompatibilities (Servedio 2004; Barton and De Cara 2009). We thus expect different evolutionary outcomes in hybrid zones. Extreme scenarios could either result in the completion of the speciation process by reinforcement or in genetic swamping and substitution of local parental genotypes by the hybrid ones (Kirkpatrick and Servedio 1999; Allendorf et al. 2001). Alternatively, hybrid zones could lead to adaptive introgression (Haines et al. 2019), formation of a new hybrid species (i.e. hybrid speciation; Hermansen et al. 2011; Lamichhaney et al. 2017), or persistence of hybrids beyond the initial generations through backcrossing and interbreeding (i.e. hybrid swarm; Abbott et al. 2013, Harrison and Larson 2014; Li et al. 2016). Finally, hybrid zones might persist over time and be sustained by the balance between selection against hybrids and continuous mating among the parental species coming from adjacent areas (i.e. tension zone model; Barton and Hewitt 1989). Consequently, the fate of a given hybrid zone relies on its ecological and genomic contexts, but our understanding of the influence of each factor is still in its infancy.

Until recently, knowledge about the mechanisms preventing populations from fully mixing and the nature of barriers to genetic exchanges relied on a few genetic markers and phenotypic variation across and within hybrid zones (e.g. Jiggins et al. 2001; Taylor et al. 2006). The advent of high-throughput sequencing has allowed us to build on this fundamental knowledge and to help characterise the underlying determinants of the observed introgression patterns and outcomes. In particular, these recent genomic approaches have provided new insight into the mosaicism of hybrid genomes and into how the genomes of the parental species are blended to form hybrid individuals (e.g. Yang et al. 2021; Zhang et al. 2023). Hybrid genomes are characterised by the presence of introgression tracts – the genetic blocks inherited from the parental species – whose size decreases with the successive generations due to recombination events. Some introgressed tracts can become fixed, while others are eliminated by random processes (i.e. genetic drift) or selection against incompatibilities (Buerkle and Rieseberg 2008). The type of selection, dominance or location on an autosome or sex chromosome influence the suppression intensity of the introgressed tracts (Hvala et al. 2018). Eventually, the interruption of genetic exchanges between the parental species and the hybrid can lead to genome stabilisation of the hybrid taxon (Buerkle and Rieseberg 2008).

During the stabilisation process, the type of selective pressure acting on the hybrid genome highly influences its architecture (Runemark et al. 2019). When hybrids mainly face ecological pressure, selection favours reproductive isolation from the parental species and adaptation to a novel niche. The genome then stabilises through the purging of incompatibilities and the maintenance of co-adapted gene complexes and pathways, resulting in a hybrid genome with both parents’ ancestry tracts (Schumer et al. 2016; Barton 2018). In the case of strong intrinsic pressure caused by hybridisation (i.e. impacting the hybrid genome integrity), selection removes the DNA of the minor parent – the species from which hybrids derive less of their genome – in gene-rich regions through backcrossing. Such mechanism will result in a stabilised genome with a few adaptive introgressed regions (Pardo-Diaz et al. 2012; Hanemaaijer et al. 2018). The proportion of the genome inherited by the hybrids from each parent might thus vary substantially within species after subsequent hybrid generations. The hybrids can exhibit mosaic genomes with an equal contribution from both parental species (e.g. Italian sparrow; Elgvin et al. 2017), a highly unbalanced contribution (*Heliconius* butterflies; Jiggins et al. 2008), or even genomes with an ancestry almost completely biased toward a single donor (e.g. *Anopheles* mosquito; Fontaine et al. 2015). Moreover, some hybrids also exhibit high variation in parental genomes proportions among individuals, as shown in some Lycaenidae butterflies, swordtail fish or Italian sparrows (Nice et al. 2013; Runemark et al. 2018a; Schumer et al. 2018). As puzzling as such patterns might be, the causes and evolutionary consequences of variable hybridisation outcomes are still poorly understood. Therefore, we need a comprehensive genomic overview of the various types of hybrid zones to draw more general conclusions and be able to predict the consequences of hybridisation on genomes.

Here, we focus on a precise location (i.e. Kimbe Bay, Papua New Guinea) of the mosaic hybrid zone (as defined by Rand and Harrison 1989) between two genetically and morphologically distinct clownfish species – *Amphiprion sandaracinos* and *Amphiprion chrysopterus*. Like all clownfish species, *A. sandaracinos* and *A. chrysopterus* exhibit a mutualistic interaction with specific host sea anemones, and have an overlap in host preference. They have a size-based hierarchy, where the largest individual in the sea anemone is the dominant female, followed in size by the male and smaller non-breeding subordinates (Fricke 1979; Buston 2003). Both species display mostly allopatric distributions but coexist in a narrow area between the Solomon Islands and the Halmahera Island (Indonesia), where they hybridise and form the hybrid *Amphiprion leucokranos* (Gainsford et al. 2015). Due to its distinctive colour patterns, the hybrid was previously considered a nominal species, but genetic, morphological and ecological data has confirmed its hybrid status (Gainsford et al. 2015). Hybrids exhibit intermediary size, colours and ecological traits compared to the parental species and, similar to the parents, use mainly *Stychodactyla mertensii* as the host sea anemone or *Heteractis crispa* as an alternative host (Gainsford et al. 2015). The size difference of parental species combined with their hierarchical social structure were suggested to be the main drivers of their mating patterns, with the larger *A. chrysopterus* being the female when reproducing with the smaller *A. sandaracinos* (Gainsford et al. 2015).

Three distinct locations (i.e. (1) Kimbe Bay and (2) Kavieng in Papua New Guinea and the (3) Solomon Islands) within the hybrid zone between *A. sandaracinos* and *A. chrysopterus* were previously studied based on a few genetic markers and morphological data (Gainsford et al. 2015; 2020). The outcome of hybridisation events in each of these locations is thought to be dependent on their specific ecological context, with the relative abundances and the distinct sizes of the parental species most likely explaining the hybridisation patterns (Gainsford et al. 2015; 2020). Mixed-species assemblages frequently occur in the Solomon Islands – where *A. chrysopterus* is mori abundant compared to *A. sandaracinos*. In addition, hybrids from both Kavieng and Solomon Islands exhibit balanced parental ancestry. In contrast, there is a bias toward *A. sandaracinos* ancestry in hybrids in Kimbe Bay, where both parental species have relatively similar frequency and mainly form conspecific assemblages (Gainsford et al. 2020). Indeed, Kimbe Bay comprises hybrids with equal contribution from both parents alongside putative *A. sandaracinos* backcrosses. However, backcrosses with *A. chrysopterus* are seldomly observed (Gainsford et al. 2015; 2020). This pattern is surprising given that hybrids inherit their mitochondrial genome from the female *A. chrysopterus* and mounting evidence that incompatible mito-nuclear interactions might result in hybrid fitness reduction (Trier et al. 2014; Hill 2016; Runemark et al. 2018a; Runemark et al. 2018b; Wagner et al. 2020).

In this study, we took a step further into understanding the dynamics of the Kimbe Bay hybrid zone location using whole-genome sequencing data. We investigated the genomic outcomes of the recurrent hybridisation among *A. sandaracinos* and *A. chrysopterus*, focusing on its impact on both the hybrids and the parental species. Given the presence of widespread hybridisation in clownfishes (Litsios et al. 2014; Marcionetti et al. 2022; Schmid et al. 2022; Marcionetti and Salamin 2023) and its potential role in their evolutionary history, we want to characterise the Kimbe Bay hybrid zone and test if it is a tension zone (Barton and Hewitt 1985) with mainly first generation hybrids or is it a hybrid swarm (Grant 1981; Allendorf et al. 2001) with a mix of early and later hybrid generations. Furthermore, we investigated the following questions: are some genomic regions more prone to be inherited by one parent or another? Are the patterns of introgression similar among hybrid individuals? What are the impacts of recurrent genetic exchanges on the parental genomes? Shedding light on these questions will improve current knowledge on the mechanisms maintaining species integrity and the hybrid genomic architecture.

## RESULTS

### Sequencing, mapping and SNP calling statistics

We achieved an average of approximately 5.91 billion raw paired reads across 67 samples, with the number of raw reads per sample ranging from ∼40 million to ∼191 million (Supp. Table S2). After trimming low-quality regions and removing low-quality reads, we ended up with approx. 36 to 173 million paired reads per sample, corresponding to an estimated coverage between 6.1X and 28.9X (Supp. Table S2).

We mapped the trimmed reads to the *A. percula* (Lehmann et al. 2018) genome and 84.9% to 89.2% of the reads mapped properly to the reference (i.e. with pairs having the correct orientation and insert-size; Supp. Table S3). We further filtered the data by removing reads mapped with low-confidence and potentially redundant sequencing data arising from the overlap of paired reads to obtain a final coverage ranging between 4.5X and 20.6X (Supp. Table S3). Following these steps, we computed genotype likelihoods, estimated the major and minor alleles and filtered the resulting VCF file to obtain a final dataset of 7,953,787 SNPs informative for the 67 samples (Supp. Table S4).

### Principal component analysis, admixture, hybrid index and mitochondrial phylogenetic reconstruction

To describe the relationship between the two parental species and the hybrid, we first performed a PCA on the covariance matrix between all individuals (Figure 1B). We found a clear separation between *A. chrysopterus* and *A. sandaracinos* – the parental species – along the first axis, which explains 91.37% of the variance. Furthermore, we highlighted two clusters of *A. leucokranos* individuals: one equally distant from both parental species clusters and the second one closer to the *A. sandaracinos* cluster. The second axis explained 2.61% of the variance and separated four *A. chrysopterus* individuals from Kimbe Bay and the Solomon Islands from the main *A. chrysopterus* cluster.

**Figure 1.**
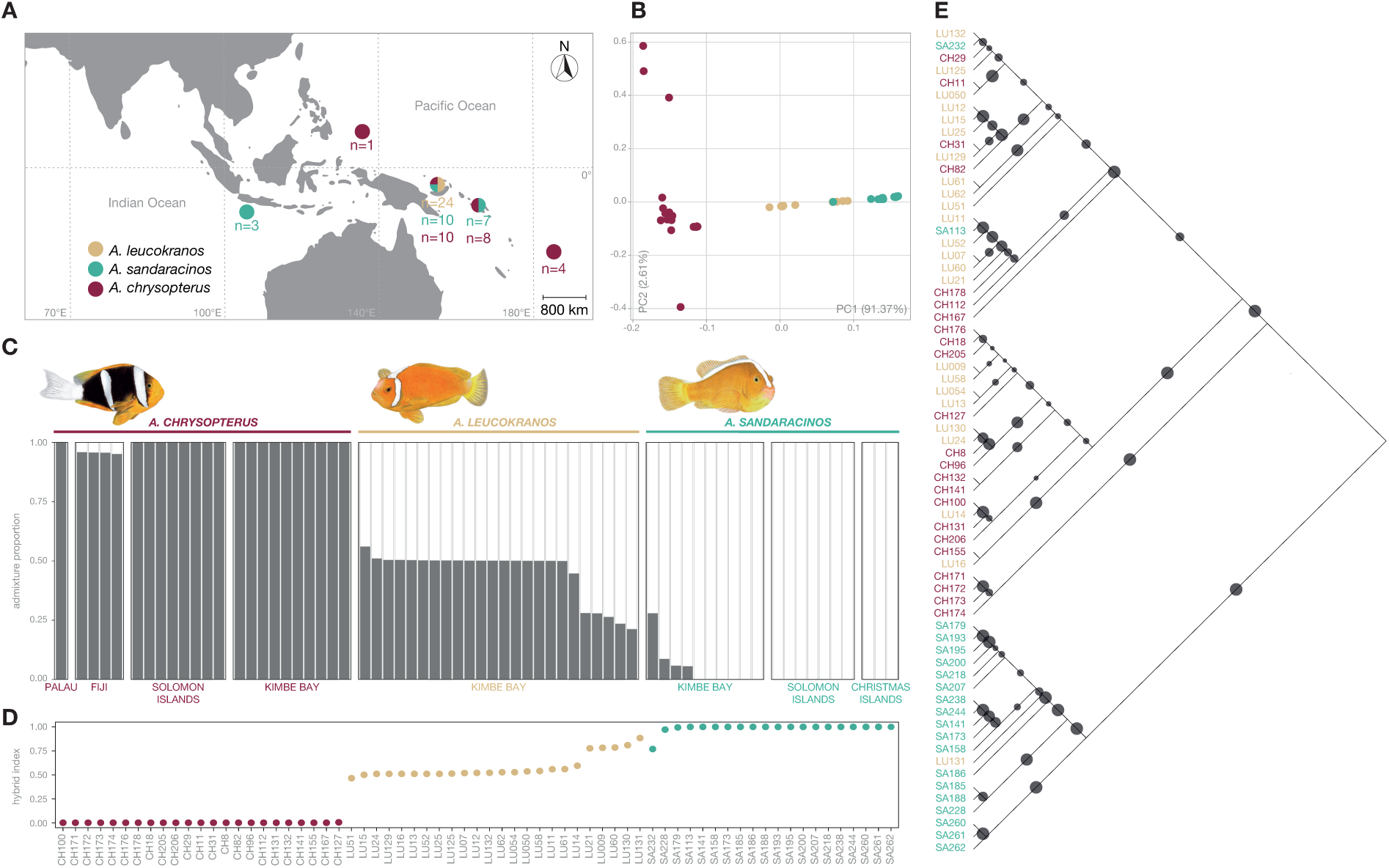
Sampling and population genetic structure. (A) Coloured circles represent sampling sites for the study. The different colours represent the three studied species: *A. chrysopterus* (dark red), *A. leucokranos* (light brown) and *A. sandaracinos* (blue). The numbers below correspond to the number of samples for each site and species. (B) Principal component analysis (PCA) showing a separation of the two parental species (*A. chrysopterus* and *A. sandaracinos*) on the first axis (PC1), with the hybrid species (*A. leucokranos*) in the middle, and separation of *A. chrysopterus* individuals from Fiji on the second axis (PC2). (C) Admixture bar plot based on NGSADMIX *q*-values for *K*=2. Each bar represents the ancestry proportion of each individual for both parental species (grey colour corresponds to *A. chrysopterus* ancestry and white to *A. sandaracinos* ancestry). (D) Hybrid index score estimated with GGHYBRID for the 67 individuals (parental species included). The colour code is the same as the PCA. (E) Mitochondrial midpoint-rooted phylogenetic reconstructed with IQ-tree V2.0.6. The phylogeny is based on the 67 mitochondrial genomes reconstructed with MITObim using *A. percula* reference mtDNA genome. The size of the dots on the branches corresponds to the bootstrap support (ranging from 11 to 100) based on 1,000 ultrafast bootstraps approximation. The colours correspond to the three species (dark red: *A. chrysopterus*; light brown: *A. leucokranos* (hybrid); turquoise: *A. sandaracinos*

We quantified the contribution of each parental species to the hybrid genome and potential introgression events in parental populations by calculating the admixture proportion of each individual (Figure 1C). The optimal number of ancestral populations (clusters) was *K* = 2 and split both parental species. At *K =* 3, *A. chrysopterus* individuals from Fiji (allopatric population) grouped in a separate cluster, and at *K* = 4, *A. sandaracinos* samples from Christmas Islands (allopatric population) formed the fourth cluster (see Figure S1 for results with *K =* 3 to *K* = 7). We found that 17 *A. leucokranos* individuals had half of each parental species ancestry (considered as F1 for subsequent analysis) and five individuals had a ∼0.75 proportion of *A. sandaracinos* ancestry (considered as backcrosses for subsequent analysis). Four *A. sandaracinos* individuals from Kimbe Bay displayed introgression from *A. chrysopterus*, the one with the highest level of admixture being probably a misidentified *A. leucokranos*.

We estimated the parental contribution of each species to the hybrid genome and checked for consistency with the admixture analysis by computing the hybrid index (HI) of each individual using a subset of informative SNPs. After filtering for LD and selecting highly differentiated SNPs among parental populations, we obtained a final dataset of 75,470 SNPs. Consistently with the admixture analysis, the hybrid index highlighted a score either close to 0.5 or close to 0.75 for all *A. leucokranos* individuals (Figure 1D). The score of the potentially misidentified *A. sandaracinos* individual was also close to 0.75.

Finally, we determined the parental inheritance of mitochondrial genome in hybrids by reconstructing a maximum-likelihood tree based on the whole mitochondrial sequence. The two methods used to reconstruct the mitochondrial genomes produced similar results (Figure 1E, Figure S2). All *A. leucokranos* clustered with *A. chrysopterus*, except for a single individual which is putatively the offspring of an *A. sandaracinos* female and a *sandaracinos*-backcross male. We also found that two *A. sandaracinos* individuals grouped with *A. chrysopterus*, one of them being the potentially misidentified and highly admixed *A. sandaracinos* (Figure 1C).

### Ancestry tracks in hybrids

To infer the local ancestry of the hybrid *A. leucokranos* individuals, we estimated allele dosage scores at each SNP using the two-layer hidden Markov model implemented in ELAI (Guan 2014). Among the 24 hybrids and consistent with the admixture analysis, 17 had entire heterozygous genomes and were subsequently considered first-generation hybrids (F1; Supp. Figure S3). One individual exhibited extended ancestry tracks from both *A. chrysopterus* and *A. sandaracinos*, a pattern consistent with a second-generation hybrid (F2; Supp. Figure S3). Finally, five individuals displayed sizable tracks of *A. sandaracinos* ancestry (similar individuals highlighted in the admixture analysis), and one had multiple blocks of *A. chrysopterus* ancestry, which is typical of early backcross generations (BC; Supp. Figure S3).

### Genomic and phenotypic characterisation of *A. leucokranos*

To explore the relationship between the 20 *A. leucokranos* individuals for which we have a picture (no picture for the four remaining hybrids), we performed two additional PCAs on these individuals only. We computed the first PCA using the whole-genome SNPs dataset (Figure 2A) and the second using the nine binary phenotypic traits characterising each hybrid (Figure 2B; see Supp. Table S5 for the matrix of phenotypes). The phenotypic traits used were based on white bands asymmetry (A), the extent of the white dorsal band (*A. sandaracinos* characteristic) and the number of vertical white stripes (*A. chrysopterus* characteristic). The first axis of the genomic PCA split the *sandaracinos*-backcross individuals from the single *chrysopterus*-backcross individual and explained 78.68% of the variance. All F1 individuals clustered together in-between the two types of backcross. The second axis explained 5.28% of the variance and isolated the dataset’s only second-generation hybrid (F2). The first and second axis of the PCA based on phenotypic data respectively explained 25.4% and 20.1% of the variance. F1 hybrids formed a cluster, whereas *sandaracinos*-backcrosses and F2 hybrids were spread across the upper-left part of the PCA space and exhibited *sandaracinos*-like phenotypic traits.

**Figure 2.**
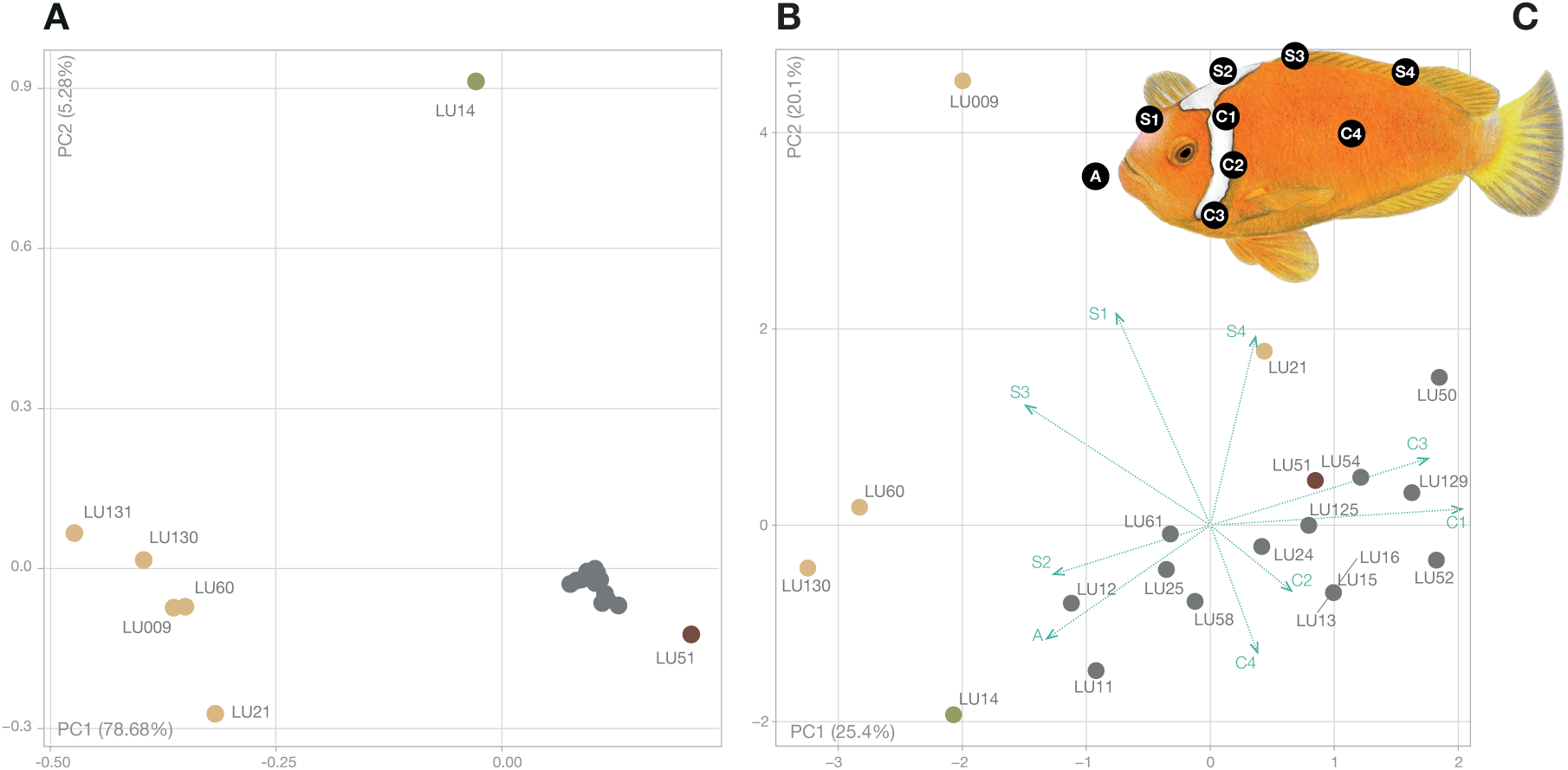
Principal component analysis (PCA) based on genetic and phenotypic data considering only *A. leucokranos* individuals. (A) PCA based on whole-genome SNPs for *A. leucokranos* only. The PC1 separates presumed *sandaracinos*-backcross individuals (light brown dots) from the *chrysopterus*-backcross individual (dark brown dot), with F1 individuals in the middle (grey dots), and PC2 separates the F2 individual (green dot) from all other hybrids. Labels correspond to the samples ID. (B) PCA based on phenotype matrix (Table S5). Blue arrows indicate loadings for each trait. The PC1 separates presumed F1 hybrids from backcross individuals, and PC2 further splits the backcross and F2 individuals. The colour code is the same as in panel (A). Labels correspond to the samples ID. (C) Illustration of the nine phenotypic characteristics used for the PCA.

### Population genomic statistics between hybrids and parental species

To investigate the genome-wide patterns of differentiation, divergence and diversity among the hybrids and the different parental species populations, we averaged the differentiation (*FST*), the absolute divergence (*dxy*) and the nucleotide diversity (*π*) values across 50 kb-genomic windows (Table 1). Pairwise *dxy* values between hybrids and the two parents were stable across the three parental populations. Comparatively, pairwise *FST* values were more variable across parental populations. They were generally lower between the hybrids and the allopatric parents (except for the comparison involving *A. sandaracinos* and backcross hybrids). Finally, nucleotide diversity was at least three times higher in both F1 and backcross hybrids compared to all parental populations.

**Table 1.** Mean *F_ST_*, *d_xy_* and nucleotide diversity (*π*) between the hybrids and the parental species.

| Parameter | Species | Parental population origins |  |  |
| --- | --- | --- | --- | --- |
|  |  | Kimbe Bay | Solomon Islands | Allopatric |
| $F_{ST}$ | CHP-BC | 0.429 ± 0.129 | 0.425 ± 0.128 | 0.329 ± 0.116 |
|  | CHP-F1 | 0.204 ± 0.036 | 0.187 ± 0.032 | 0.136 ± 0.022 |
|  | SAN-BC | 0.102 ± 0.068 | 0.115 ± 0.073 | 0.113 ± 0.049 |
|  | SAN-F1 | 0.165 ± 0.029 | 0.183 ± 0.033 | 0.106 ± 0.021 |
|  | BC-F1 | 0.016 ± 0.031 | - | - |
| $d_{xy}$ | CHP-BC | 0.577 ± 0.087 | 0.576 ± 0.087 | 0.537 ± 0.068 |
|  | CHP-F1 | 0.442 ± 0.028 | 0.443 ± 0.028 | 0.443 ± 0.028 |
|  | SAN-BC | 0.275 ± 0.079 | 0.276 ± 0.079 | 0.298 ± 0.070 |
|  | SAN-F1 | 0.419 ± 0.033 | 0.420 ± 0.034 | 0.421 ± 0.032 |
|  | BC-F1 | 0.426 ± 0.029 | - | - |
| $\pi$ | CHP | 0.107 ± 0.050 | 0.106 ± 0.050 | 0.087 ± 0.043 |
|  | SAN | 0.064 ± 0.055 | 0.060 ± 0.050 | 0.066 ± 0.046 |
|  | BC | 0.376 ± 0.074 | - | - |
|  | F1 | 0.442 ± 0.028 | - | - |
| Allopatric corresponds to the *A. chrysopterus* Fiji population and *A. sandaracinos* Christmas Islands population. All comparisons and populations statistics are in Supp. Table S6. CHP: *A. chrysopterus*; SAN: *A. sandaracinos* (admixed individuals removed); BC: backcross with *A. sandaracinos*; F1: first-generation hybrids. |  |  |  |  |

### Characterisation of backcross’ genomic architecture

Through the admixture and local ancestry analyses, we detected five *A. sandaracinos* backcross individuals and we characterised in these individuals the genomic tracts inherited from *A. sandaracinos*. The distribution of tracks with *A. sandaracinos* ancestry in backcross individuals was heterogeneous across the genome. Most chromosomes displayed introgressed tracks in all backcrosses, whereas chromosome 8 had no *A. sandaracinos* introgressed regions (Figure 3). A percentage of 21.3% of 50kb-windows (i.e. 3,617 windows) displayed *A. sandaracinos* ancestry in at least four out of the five backcross individuals. All *FST*, *dxy*, and *π* (except for the *dxy* between *A. chrysopterus* and F1-hybrids) were significantly different between the introgressed tracts and the background genomic windows (Table 2). However, the effect size was negligible or small for most comparisons. The only exceptions are the comparisons involving statistics related to backcross hybrids, for which we detected a large effect size for the differences in nucleotide diversity and both divergence and differentiation involving the two parental species (Table 2). Furthermore, the observed frequency of putative barrier loci (see material and methods for definition) was not different from the expected frequency (*χ*^2^ = 3, *p*-value = 0.08; Table 2). Finally, we ran a gene ontology analysis on the 4,734 genes located in the introgressed windows and found 30 significantly enriched terms (*p*-value < 0.01; Supp. Table S7). Among the 30 significant GO terms, twelve GO were associated with locomotion (e.g. GO:0040013, GO:0048521) and seven with the regulation of transferase activity (e.g. GO:0051443, GO:0045745). One GO term was also associated with the regulation of biological processes involved in symbiotic interaction (GO:0043903).

**Figure 3.**
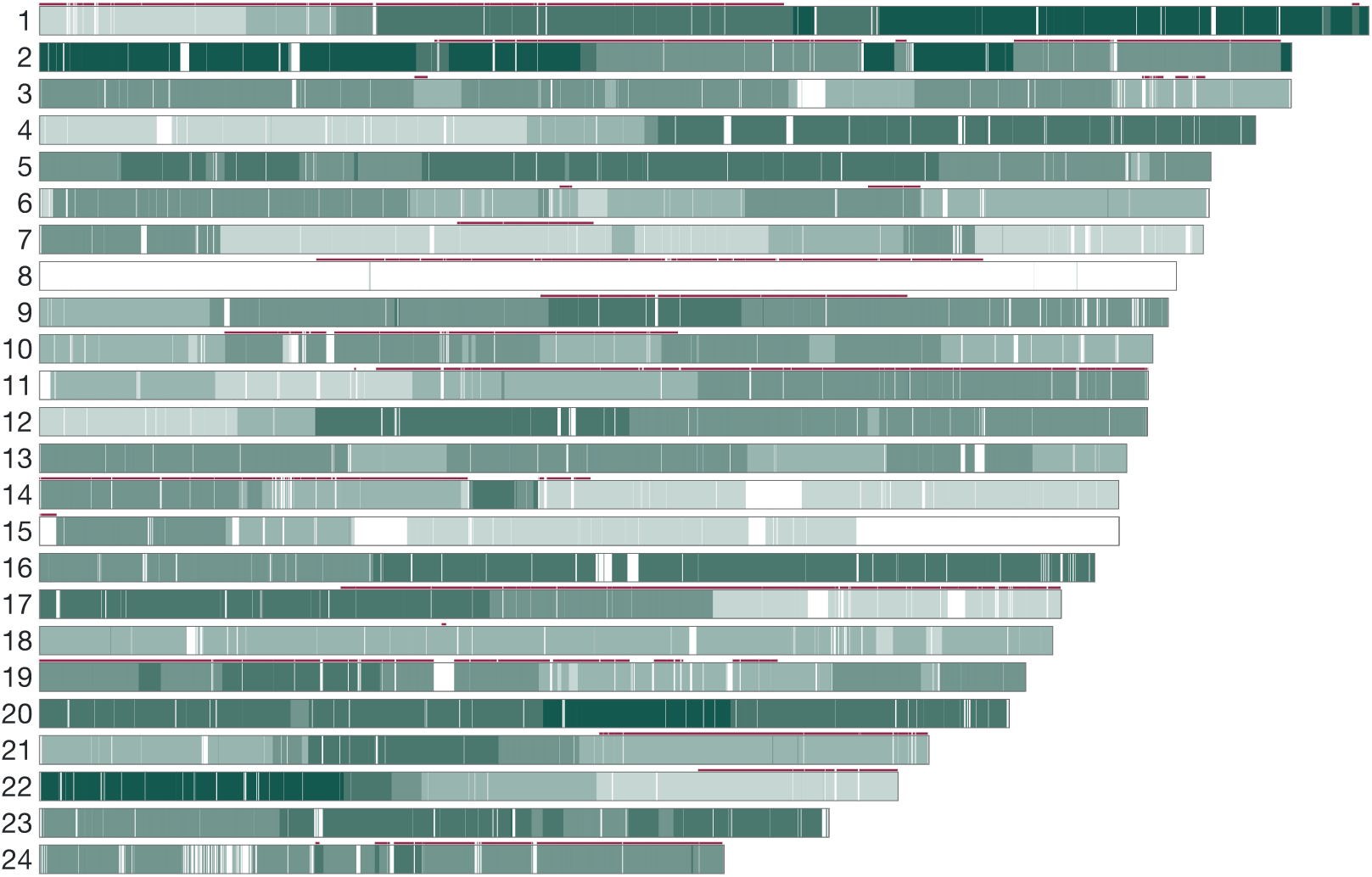
Karyotype of the parental ancestry estimated with ELAI (Guan 2014) across the 24 chromosomes of backcross *A. leucokranos*. The shade of green corresponds to the number of backcrosses with *A. sandaracinos* ancestry and ranges from zero individuals (white) to five individuals (dark green). The red line above each chromosome highlights regions with *A. chrysopterus* ancestry in the only *A. chrysopterus*-backcross individual.

**Table 2.**
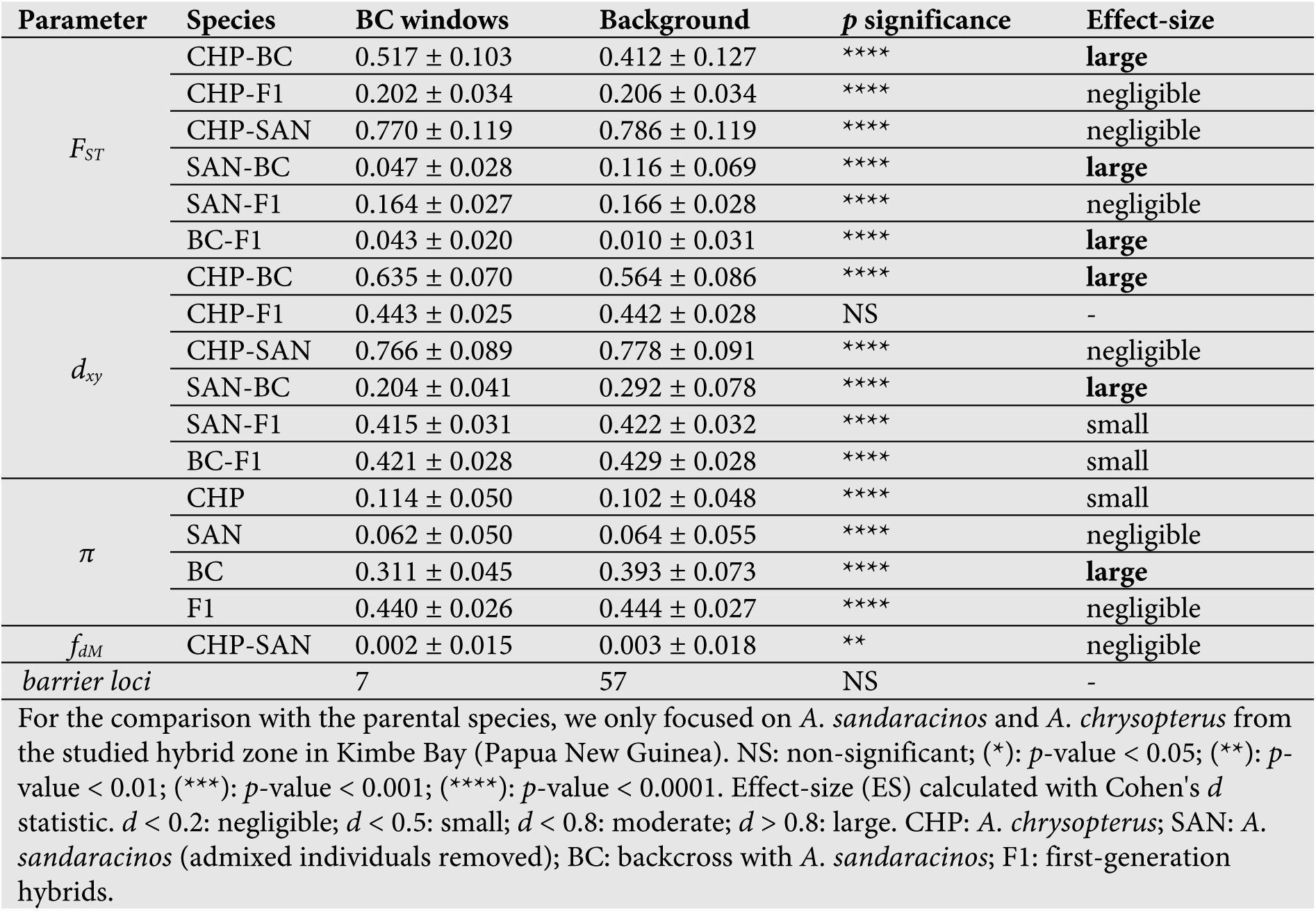
Mean population genomics statistics and number of putative barrier loci for the introgressed *A. sandaracinos* windows in the backrcross individuals compared to the genomic background.

### Genomic landscape of F1-hybrids

The admixture and local ancestry analyses highlighted that 17 *A. leucokranos* were first-generation (F1) hybrids between *A. sandaracinos* and *A. chrysopterus*. To investigate the relationship between the F1-hybrids and the parental species in sympatry (Kimbe Bay), we compared the level of differentiation (*FST*) between the hybrids and both parents to identify highly divergent regions of the genome. The distribution of the *FST* difference across the genome was significantly biased towards higher differentiation between the F1-hybrid and *A. chrysopterus* (mean difference significantly different from 0; *p*-value < 2E-16; Figure 4A).

**Figure 4.**
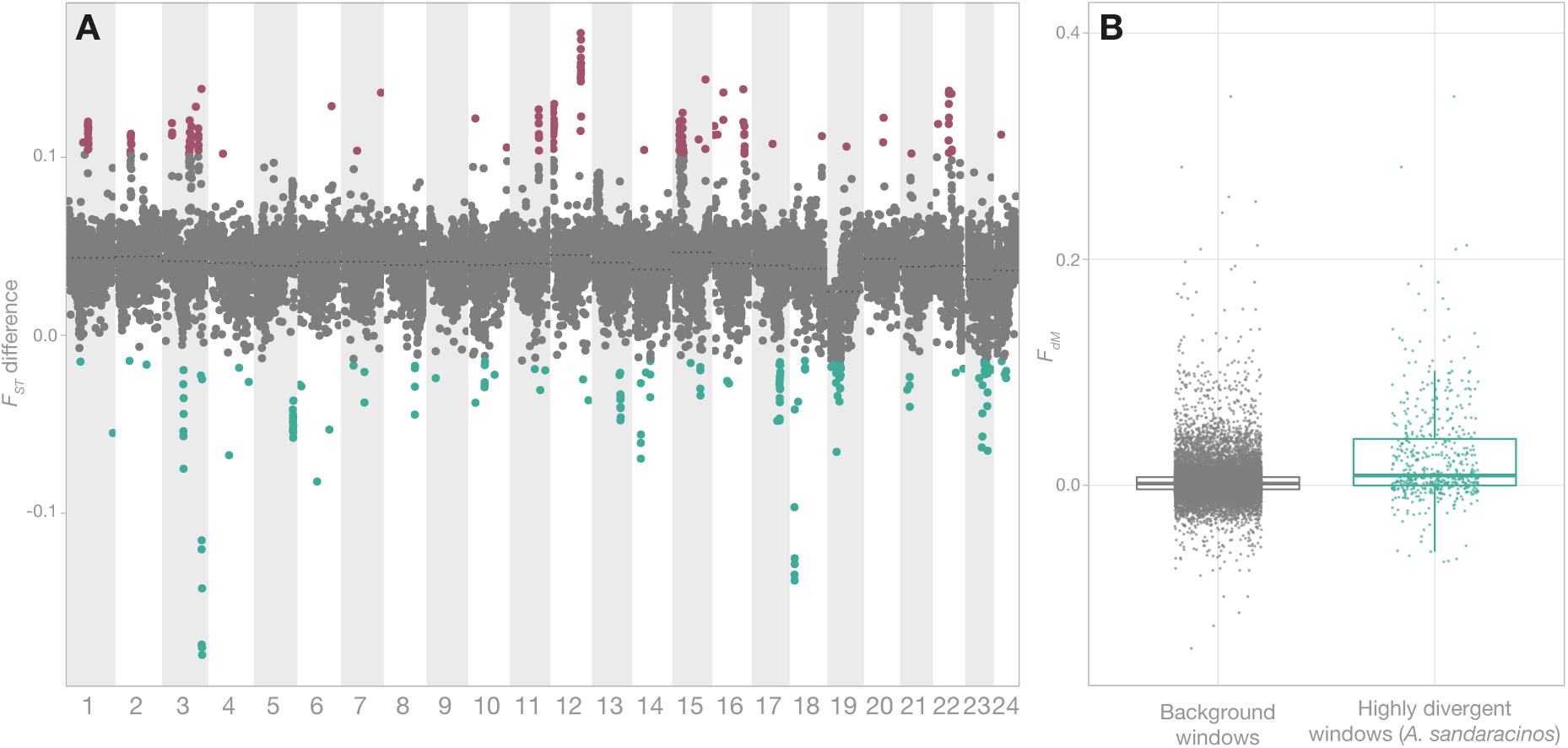
Characterisation of the F1-hybrids genomic landscape. (A) *F_ST_* difference between F1-hybrids and both parental species from Kimbe Bay (CHP-F1 *F_ST_* – SAN-F1 *F_ST_*). Red points correspond to values above the 99^th^ percentile (high *F_ST_* for *A. chrysopterus*-F1 and low *F_ST_* for *A. sandaracinos*-F1), and blue points highlight values below the 1st percentile (low *F_ST_* for *A. chrysopterus*-F1 and high *F_ST_* for *A. sandaracinos*-F1). Both colored points were considered highly divergent windows. The mean *F_ST_* difference for each chromosome is indicated with a dotted line. (B) Distribution of *f_dM_* across all background windows (grey) and across the highly divergent windows biased toward *A. sandaracinos* (blue). Mean difference was highly significant (*p*-value = 4.4e-43) and had a moderate effect size (*d* = - 0.65)

The number of highly divergent windows per chromosome was not statistically different from the expected distribution generated with 10,000 permutations for most chromosomes, except for chromosomes 3, 12, 15 and 19, which exhibited an inflated number of outliers (i.e., respectively 40, 34, 47 and 25 windows; Figure 4A). Although all population genomic statistics were statistically different between the background and highly divergent windows, the effect size was moderate or large in only part of the statistics (Table 3). Notably, *A. sandaracinos* exhibited higher nucleotide diversity in divergent windows below the 1^st^ percentile compared to the background windows. Furthermore, the *fdM* statistic was higher in the divergent windows showing a bias towards *A. sandaracinos* compared to the background windows (Figure 4B). Finally, we identified 609 genes in the highly divergent windows and found 21 significant GO terms (*p*-value < 0.01; Supp. Table S8). Among them, 14 were associated with various processes linked to muscles (e.g., GO:0003300, GO:0007274), and four were related to peptidyl-tyrosine phosphorylation (GO:0018108, GO:0006283, GO:0006164, GO:0000462).

**Table 3.**
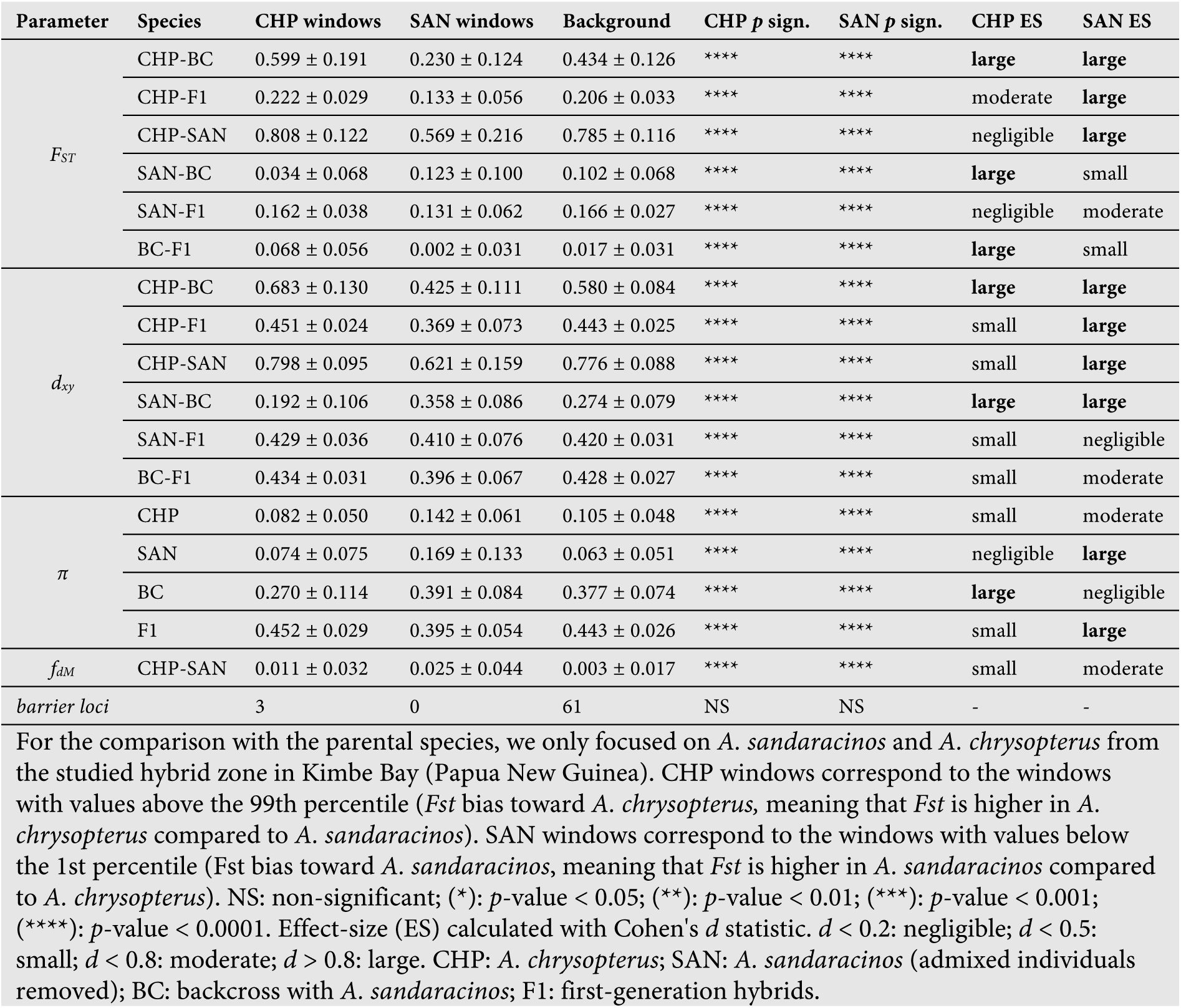
Mean population genomics statistics for highly divergent windows between F1-hybrids and parental species.

| Parameter | Species | CHP windows | SAN windows | Background | CHP *p* sign. | SAN *p* sign. | CHP ES | SAN ES |
| --- | --- | --- | --- | --- | --- | --- | --- | --- |
| $F_{ST}$ | CHP-BC | 0.599 ± 0.191 | 0.230 ± 0.124 | 0.434 ± 0.126 | **** | **** | large | large |
|  | CHP-F1 | 0.222 ± 0.029 | 0.133 ± 0.056 | 0.206 ± 0.033 | **** | **** | moderate | large |
|  | CHP-SAN | 0.808 ± 0.122 | 0.569 ± 0.216 | 0.785 ± 0.116 | **** | **** | negligible | large |
|  | SAN-BC | 0.034 ± 0.068 | 0.123 ± 0.100 | 0.102 ± 0.068 | **** | **** | large | small |
|  | SAN-F1 | 0.162 ± 0.038 | 0.131 ± 0.062 | 0.166 ± 0.027 | **** | **** | negligible | moderate |
|  | BC-F1 | 0.068 ± 0.056 | 0.002 ± 0.031 | 0.017 ± 0.031 | **** | **** | large | small |
| $d_{xy}$ | CHP-BC | 0.683 ± 0.130 | 0.425 ± 0.111 | 0.580 ± 0.084 | **** | **** | large | large |
|  | CHP-F1 | 0.451 ± 0.024 | 0.369 ± 0.073 | 0.443 ± 0.025 | **** | **** | small | large |
|  | CHP-SAN | 0.798 ± 0.095 | 0.621 ± 0.159 | 0.776 ± 0.088 | **** | **** | small | large |
|  | SAN-BC | 0.192 ± 0.106 | 0.358 ± 0.086 | 0.274 ± 0.079 | **** | **** | large | large |
|  | SAN-F1 | 0.429 ± 0.036 | 0.410 ± 0.076 | 0.420 ± 0.031 | **** | **** | small | negligible |
|  | BC-F1 | 0.434 ± 0.031 | 0.396 ± 0.067 | 0.428 ± 0.027 | **** | **** | small | moderate |
| $\pi$ | CHP | 0.082 ± 0.050 | 0.142 ± 0.061 | 0.105 ± 0.048 | **** | **** | small | moderate |
|  | SAN | 0.074 ± 0.075 | 0.169 ± 0.133 | 0.063 ± 0.051 | **** | **** | negligible | large |
|  | BC | 0.270 ± 0.114 | 0.391 ± 0.084 | 0.377 ± 0.074 | **** | **** | large | negligible |
|  | F1 | 0.452 ± 0.029 | 0.395 ± 0.054 | 0.443 ± 0.026 | **** | **** | small | large |
| $f_{dM}$ | CHP-SAN | 0.011 ± 0.032 | 0.025 ± 0.044 | 0.003 ± 0.017 | **** | **** | small | moderate |
| barrier loci |  | 3 | 0 | 61 | NS | NS | - | - |

For the comparison with the parental species, we only focused on *A. sandaracinos* and *A. chrysopterus* from the studied hybrid zone in Kimbe Bay (Papua New Guinea). CHP windows correspond to the windows with values above the 99th percentile (*Fst* bias toward *A. chrysopterus*, meaning that *Fst* is higher in *A. chrysopterus* compared to *A. sandaracinos*). SAN windows correspond to the windows with values below the 1st percentile (*Fst* bias toward *A. sandaracinos*, meaning that *Fst* is higher in *A. sandaracinos* compared to *A. chrysopterus*). NS: non-significant; (\*): *p*-value < 0.05; (\*\*): *p*-value < 0.01; (\*\*\*): *p*-value < 0.001; (\*\*\*\*): *p*-value < 0.0001. Effect-size (ES) calculated with Cohen's *d* statistic. *d* < 0.2: negligible; *d* < 0.5: small; *d* < 0.8: moderate; *d* > 0.8: large. CHP: *A. chrysopterus*; SAN: *A. sandaracinos* (admixed individuals removed); BC: backcross with *A. sandaracinos*; F1: first-generation hybrids.

### Population genomics of the parental species in the hybrid zone and outside

Having confirmed the occurrence of early generation *A. leucokranos* hybrids (i.e F1, F2 and backcrosses), we conducted population genomic analyses of the two parental species in Kimbe Bay to explore the impact of ongoing genetic exchanges and recurrent backcrossing in *A. sandaracinos* (Figure 5A). Mean differentiation and divergence across the genome between *A. sandaracinos* and *A. chrysopterus* were respectively 0.780 (± 0.125) and 0.773 (± 0.09). The mean nucleotide diversity was higher in *A. chrysopterus* (0.101 ± 0.05) compared to *A. sandaracinos*(0.065 ± 0.056). Finally, although the mean *fdM* value across the genome was close to zero (0.003), we found some positive peaks in chromosomes 3, 12, 15, 20 and 21 and a single negative peak in chromosome 19 (Figure 5A).

**Figure 5.**
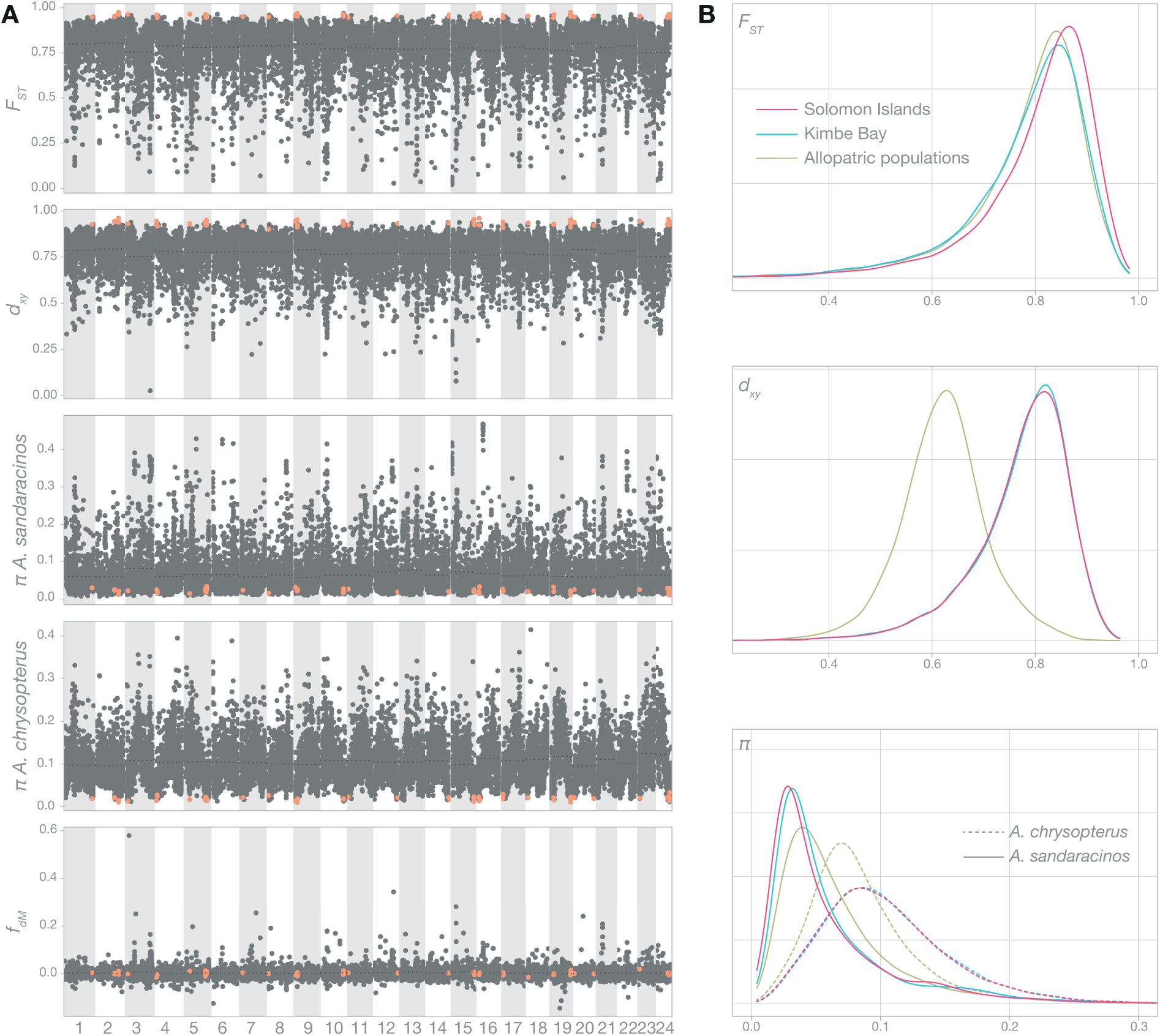
Genomic landscape of parental species in the hybrid zone and compared to other populations. (A) *F_ST_*, *d_xy_*, *f_dM_*, and *π* in 50kb-windows across the genome of *A. sandaracinos* and *A. chrysopterus* in Kimbe Bay. Orange points correspond to potential barrier loci identified using outlier *F_ST_* values among the three parental populations. Mean values for each chromosome are indicated with a dotted line. (B) Density plot of *F_ST_*, *d_xy_*, and *π* for Kimbe Bay (blue), Solomon Islands (pink) and for two populations (Fiji and Christmas Islands) where the two parental species are in allopatry (green). *F_ST_* and *d_xy_* were computed between *A. sandaracinos* and *A. chrysopterus* occurring in Kimbe Bay, between *A. sandaracinos* and *A. chrysopterus* occurring in Solomon Islands and between *A. sandaracinos* from Christmas Islands and *A. chrysopterus* from Fiji to compare populations where the two species are in allopatry.

We also compared the density distribution of the same population genomics statistics to other *A. sandaracinos*-*A. chrysopterus* pairwise populations either in sympatry (Solomon Islands) or in allopatry (Fiji and Christmas Islands; Figure 5B). Although the *FST* distributions were similar among the three pairwise population comparisons, the *dxy* distribution of the allopatric populations was shifted toward smaller values compared to the two sympatric populations. Finally, nucleotide diversity had highly similar distributions for both species in Kimbe Bay and the Solomon Islands. However, the allopatric *A. sandaracinos* displayed higher nucleotidic diversity compared to the two sympatric populations, whereas the allopatric *A. chrysopterus* population exhibited a distribution that was shifted toward lower values (Figure 5B, see Supp. Figure S4 for all pairwise comparisons).

## DISCUSSION

Hybrid zones are robust systems for studying the interaction between diverging gene pools and thus provide the basis for understanding the mechanisms leading to speciation (Barton and Hewitt 1985; Gompert and Buerkle 2016). Besides their importance in studying evolutionary processes, hybrid zones are puzzling phenomena displaying their own evolutionary history, as their fate ranges from population collapse to generation of wide admixture zones (Abbott et al. 2013; Arnold 1997). Here, we conducted whole-genome analyses of the Kimbe Bay hybrid zone between two clownfish species, *A. sandaracinos* and *A. chrysopterus*. We aimed to investigate the impact of hybridisation on the integrity of the parental species and characterise the genomic architecture of the resulting hybrid *A. leucokranos.* Our analyses showed that besides the parental species, the hybrid zone comprised first-generation hybrids (F1) and early backcross generations with *A. sandaracinos*, whereas later generation hybrids and backcrosses with *A. chrysopterus* were scarce. An in-depth investigation of the genomic architecture of the backcross individuals revealed that the introgression of *A. sandaracinos* into the hybrid genome was unbalanced between chromosomes. The genomic examination of the parental species integrity highlighted high-level introgression concentrated in some genomic regions and a potential flow of *A. chrysopterus* genome into *A. sandaracinos* through the hybrids. The Kimbe Bay hybrid zone might thus act as a conduit for transferring potentially adaptive alleles from one parental species to the other.

### Hybrid zone population structure

Based on whole-genome analysis of the Kimbe Bay hybrid zone, we found that the two parental species were mainly co-occurring with first-generation (F1) hybrids and early-generation *A. sandaracinos*-backcrosses (Figure 1, Figure 2). Such hybrid composition does not match the typical hybrid zone categorisations suggested by Jiggins and Mallet (2000), which aims at evaluating the level of reproductive isolation among the parental lineages.

This characterisation typically relies on the relative composition of hybrids and parents. Bimodal hybrid zones mainly constituted by the parental forms involve strong reproductive isolation between the parents, whereas unimodal hybrid zones – consisting in a mix between various hybrid generations and the parents (i.e. hybrid swarm) – are caused by weak reproductive barriers (Jiggins and Mallet 2000). Similar to pufferfishes (Takahashi et al. 2017) and North Atlantic eels hybrid zones (Pujolar et al. 2014), the Kimbe Bay hybrid zone coincides more with a trimodal hybrid zone; parents and early generation hybrids coexist (Gay et al. 2008), even though the presence of early-generation *A. sandaracinos* backcrosses is unexpected under this model. This asymmetry in introgression was previously attributed to various factors, including differences in generation time (Barton 1986), mating behaviour (Konkle and Philipp 1992; Lamb and Avise 1986), fitness (Ostberg et al. 2013) or relative abundances of parental species (Lepais et al. 2009). In the Kimbe Bay hybrid zone, the abundance and generation time of parental species are comparable (Gainsford et al. 2020) and are unlikely to generate the observed pattern. However, as Gainsford et al. (2020) suggested, this could be a consequence of the size-based hierarchy of the clownfish. Indeed, similar to numerous fish species, reproductive success is biased toward dominant individuals, which are usually bigger and more aggressive, and thus benefit from a priority for reproduction and other limiting resources (Ang and Manica 2010; Keller and Reeve 1994; Wong 2011). As a result, the intermediary-sized hybrids might choose to reproduce with the smaller-sized *A. sandaracinos*, since the hybrids are usually ignored by the bigger *A. chrysopterus*, which favour mating with conspecific individuals (Gainsford et al. 2020). Such mechanisms are further supported by the observation that heterospecific assemblages of *A. sandaracinos*-hybrids were more common in Kimbe Bay compared to *A. chrysopterus-*hybrid or only hybrids groups (Gainsford et al. 2020).

We also found that the hybrid zone was characterised by a paucity of late-generation hybrids (i.e. *A. leucokranos* x *A. leucokranos*). Indeed, the frequency of F1-hybrids and the prevalence of backcrossing with *A. sandaracinos* confirm that the hybrids are fertile and viable. In addition, the mosaic structure of the Kimbe Bay hybrid zone – which displays various combinations of sea anemone host species, depth and substrate – should be well suited for the formation of a hybrid swarm, which is defined as a mix between early and late hybrid generations (Grant 1981; Allendorf et al. 2001). The lack of late-generation hybrids might therefore be a consequence of their reduced fitness, as previously observed in numerous studies (Arnegard et al. 2014; Christie and Strauss 2018; Orr 1995; Svedin et al. 2008). Different mechanisms could explain this pattern. One prominent explanation is selection against hybrids due to Dobzhansky-Muller incompatibilities (DMIs; Orr 1995). Indeed DMIs are increasingly revealed after the first F1 hybrid generation due to the combined effect of recombination and independent assortment, which breaks down gene complexes that coevolved over thousands of generations (Burton et al. 2006; Edmands and Burton 1999; Ellison and Burton 2008; Stebbins 1959). This process – known as “hybrid breakdown” – usually has a more severe impact on F2 hybrids compared to F1 hybrids and was highlighted in various hybridising fish species (Bolnick and Near 2005; Stelkens et al. 2015; Carlon et al. 2021; McKenzie et al. 2021; Šimková et al. 2022). In addition, the intermediate phenotypes of hybrids can be suboptimal in an environment where both parental species display phenotypic optimum for their specific niche (Barton 2001; Simon et al. 2018; Thompson 2020). In the clownfish-specific case, this can be translated into the availability of host sea anemone, which is a limiting resource (Litsios et al. 2014). The hybrids do not display transgressive phenotypes enabling them to colonise distinct host anemone species compared to the parental species (Gainsford et al. 2015). Thus, they rarely form conspecific assemblages and predominantly share the host sea anemone with one of the parental species (Gainsford et al. 2020), which might prevent them from mating with other hybrids.

The phenotypically identified hybrids were mainly F1 or first-generation backcrosses with *A. sandaracinos*. However, some individuals identified as *A. sandaracinos* based on their phenotype displayed a substantial amount of introgression with *A. chrysopterus* (Figure 1) and might consist in descendants of multiple generations of backcross with *A. sandaracinos*. Thus, only a few generations of backcrossing appear to erase *A. chrysopterus* phenotypic characteristics, resulting in backcross individuals identical to the pure parental species, a pattern described in a sparrow hybrid zone (Walsh et al. 2015). If such a mechanism is at play, it might have biased our sampling, which relies on the individuals’ phenotype. Indeed, by selecting putative hybrids based on their phenotype, we probably missed backcross individuals which are phenotypically totally similar to *A. sandaracinos*, thus biasing our sampling towards F1-generation hybrids. Therefore, analysis of additional *A. sandaracinos* coming from the Kimbe Bay hybrid zone would be required to evaluate the true extent of backcrossing.

### Hybrid zone genomic architecture and potential outcomes

We detected considerable introgression variation across the genome of backcross individuals (Figure 3). Indeed, some regions were consistently introgressed from *A. sandaracinos* in most backcrosses (notably in chromosomes 1 and 2), whereas other chromosome segments displayed a total lack of introgression. Such results are expected under the semipermeable view of species boundaries, which translate into differential introgression patterns across the genome (Harrison 1990). However, the introgressed genomic windows did not present dissimilar differentiation, divergence and nucleotide diversity values compared to the background windows (Table 2), which might have hinted at the potential selection forces at play. Such lack of difference might result from the length of the introgressed genomic tracts in the backcrosses. Indeed, the backcross individuals in our dataset correspond to early backcross generations and thus exhibit large tracts of DNA inherited by the parental species. The low number of generations implies that recombination did not have time yet to break down the large introgressed tracts into smaller genomic regions, thus preventing the detection of loci that may have been selected and fixed (Navascués et al. 2020). Despite this lack of resolution to detect selection signals in the introgressed tracts and the limited number of backcross samples, the apparent absence of introgression in chromosome 8 is intriguing. Loci under divergent selection (and potentially linked regions) associated with population or species divergence usually present particularly reduced introgression rates (Barton and Hewitt 1985; Payseur 2010; Harrison and Larson 2014). Interestingly *casz1* and various *hoxc* genes are located on this chromosome and were strongly associated with colour patterns in the hamlet reef fishes (*Hypoplecturus* spp; Hench et al. 2022). This chromosome might thus have an essential role in determining the hybrid colour pattern. However, such a conclusion is highly speculative, and further investigations are required to link the absence of introgression and the presence of colouration genes.

Although we did not detect any evolutionary relevant signal in the introgressed tracts, features linked with the genomic landscape likely contributed to the pattern of introgression in clownfish species. Indeed, an increasing amount of studies demonstrated that introgression rate is variable across the genome of many species (e.g. Janoušek et al. 2015, Kovach et al. 2016, Rafati et al. 2018, Hagberg et al. 2022), including fish (e.g. Nolte et al. 2009; Schaefer et al. 2016). Beyond the well-known case of sex chromosomes subject to reduced introgression (Presgraves 2008; Maheshwari and Barbash 2011; Burton and Barreto 2012; Trier et al. 2014) – not particularly relevant for species displaying a socially-controlled sex change such as clownfish – some genomic patterns common to multiple independent hybridisation events start to emerge. Notably, genomic regions exhibiting a reduced density of coding genes or conserved elements experience an increased level of introgression (Brandvain et al. 2014; Sankararaman et al. 2016; Schumer et al. 2018; Martin et al. 2019). In addition, introgression is favoured in the genomic portions subject to an elevated recombination rate (Sankararaman et al. 2014; Janoušek et al. 2015; Sankararaman et al. 2016). This correlation results from recombination breaking down the association between neutral or adaptive alleles and deleterious alleles, eventually decreasing the strength of selection against the introgressed tracts (Schumer et al. 2018; Martin et al. 2019; Calfee et al. 2021; Veller et al. 2021). Genomic architecture can thus impact the level of introgression and partially predict the outcome of hybridisation events (Brandvain et al. 2014; Ravinet et al. 2018; Schumer et al. 2018; Martin et al. 2019; Nelson et al. 2020). Although we are missing crucial information such as the recombination rate across the genome to predict the Kimbe Bay hybrid zone outcomes accurately, we can still formulate some hypotheses based on the observed patterns of introgression and population structure of the hybrid zone.

The coexistence of early-generation *A. sandaracinos*-backcrosses with *A. sandaracinos* individuals exhibiting low-level of admixture (see Figure 1C) suggests that hybrid genome stabilisation could occur through multiple backcross generations and subsequent reduction of the minor parent genome (*A. chrysopterus*). The resulting stabilised hybrid genome would consist of a few potentially adaptive introgressed tracts of *A. chrysopterus* embedded in an *A. sandaracinos* background (Runemark et al. 2019), similar to the stabilised hybrid genomes of the *Anopheles* mosquito (Hanemaaijer et al. 2018) or the monkeyflowers (Brandvain et al. 2014). This hypothetical outcome is further supported by the few peaks of introgression we detected between the *A. sandaracinos* and *A. chrysopterus* in the hybrid zone, suggesting that past genetic exchanges occurred between the parental species (Figure 5). However, the time to reach genome stabilisation is difficult to predict since high variability exists among species. For instance, fixation of ancestry blocks was swift in experimental *Helianthus* sunflower hybrids (Rieseberg et al. 1996) and introgressed Neanderthal regions became fixed in the human genome ca. 2’000 generations following hybridisation (Sankararaman et al. 2014).

Another hybrid zone outcome highlighted in various species is hybrid speciation (Hermansen et al. 2011; Mavárez et al. 2006; Rieseberg et al. 2003). Homoploid hybrid speciation requires reproductive isolation of the hybrid against both parental species, which can arise through reproductive barriers acting before or after fertilisation (Abbott et al. 2013; Schumer et al. 2014a). Assortative mating between hybrids (Melo et al. 2009; Selz et al. 2014; Lamichhaney et al. 2017), genomic structural disparities (Rieseberg 2001; Lai et al. 2005), sorting of parental incompatibilities (Hermansen et al. 2014) or adaptation to novel ecological niches through transgressive phenotypes (Schwarzbach et al. 2001) are among the multiple barriers that could lead to homoploid hybrid speciation. However, despite one putative F2 hybrid in our dataset, we have found no evidence of ongoing mating between hybrids, suggesting that homoploid hybrid speciation is an unlikely scenario. The Kimbe Bay hybrid zone might thus promote the exchange of adaptive alleles among the parental species through the hybrid individuals.

### Parental species: ongoing genetic exchanges, impact and comparison with other populations

We detected high differentiation and divergence throughout the genome of the parental species *A. sandaracinos* and *A. chrysopterus* (Figure 5). Such divergence was expected since the two species shared a common ancestor ca. 7.5 MYA – a considerable evolutionary time considering that the clownfish radiation started ca. 14 MYA. Nevertheless, the occurrence of F1-generation and early backcross hybrids in the Kimbe Bay hybrid zone (Figure 1) indicates ongoing genetic exchanges between the two parental species and confirms that a high level of genetic divergence is a poor predictor of the ability to hybridise (Jiggins and Mallet 2000). The ongoing genetic exchange between the two parental species is not surprising since clownfishes are known for recurrent past hybridisation (Schmid et al. 2022; Marcionetti and Salamin 2023) and frequently hybridise when raised in aquaria (Fautin and Allen 1997). Furthermore, the shared use of resources – here, the host sea anemone species shared by hybridising individuals – might also favour hybridisation, as for other reef fish species (Montanari et al. 2016). Reproductive isolation is thus not complete yet, raising the question of the barriers enabling the two parental species to maintain their integrity in the face of genetic exchange. Conflict in the genomes of the parental species (i.e. genetic incompatibilities) appear to maintain species boundaries in a swordfish hybrid zone despite frequent interbreeding and overlapping environments (Schumer et al. 2014b). In addition, mating behaviour and reduced fitness of second-generation hybrids due to the accumulation of DMIs were suggested to preserve intact species boundaries in parrotfishes (Carlon et al. 2021). Beyond these examples in fishes, other species exhibit various prezygotic (Poelstra et al. 2014; Sobel et al. 2019; Stankowski et al. 2015; Toews et al. 2016) and postzygotic barriers (Brelsford and Irwin 2009; Singhal and Moritz 2012; Pulido-Santacruz et al. 2018) which maintain parental species integrity if sufficiently strong. However, genetic exchange might be able to dissolve those barriers provided the latter are weak enough (Barton and Bengtsson 1986; Gavrilets 1997; Kondrashov 2003; Xiong and Mallet 2022).

In the Kimbe Bay hybrid zone, genetic exchange seems to impact the integrity of one of the parental species. Indeed, the difference in pairwise *FST* among F1-hybrids and both parental lineages exhibit a unexpected biased distribution, with most genomic windows revealing a lower differentiation between the F1-hybrids and *A. sandaracinos*compared to *A. chrysopterus* (Figure 4A). Since the parental species are highly differentiated (Table 1, Figure 5), a large proportion of SNPs should be alternatively fixed in each lineage (i.e. homozygous for the reference allele in one parent and homozygous for the alternative allele in the other). The remaining SNPs are thus expected to be heterozygous in some individuals. They should be randomly distributed across the genomes of the parental species, generating an expected distribution of *FST* differences between the F1-hybrid and both parents centred around zero. The unexpected pattern we observed is most likely a consequence of the genomic features of the parental species rather than a process impacting F1-hybrids directly since the latter should be shielded against intrinsic incompatibilities because hybrid breakdown usually emerges in the second generation (Dobzhansky 1950; Endler 1977; Burton 1990). A potential explanation could be that genetic exchange among the parental species alters the *A. sandaracinos* genomic integrity. The recurrent unidirectional genetic exchange due to backcrossing might thus impact *A. sandaracinos* population allelic frequencies, which might start to be more similar to the ones from *A. chrysopterus*. Consequently, F1-hybrids – which are expected to be totally heterozygous – would display a reduced whole-genome differentiation against *A. sandaracinos* compared to the differentiation against *A. chrysopterus*, leading to the observed bias. As a consequence of the unilateral introgression via hybrids due to size dimorphism in parental species, we might thus witness the diffusion of *A. chrysopterus* loci into *A. sandaracinos* in the Kimbe Bay hybrid zone.

Furthermore, outlier windows strongly biased towards one parent species or the other displayed some genomic statistics that differed from the background windows (Table 3). Among them, the mean *fdM* values – which described introgression among the parental species – were higher in outlier windows compared to the background, with a larger effect size when only considering the highly differentiated windows between *A. sandaracinos* and the F1-hybrids but weakly differentiated between *A. chrysopterus* and the F1-hybrids (Figure 4B). The outlier windows are thus related to past genetic exchange among the parental species and might reflect regions of low divergence between the parents due to the fixation of introgressed alleles. Past and ongoing genetic exchange thus shape the parental species genome, as highlighted in multiple fish hybrid zones (e.g. Keller et al. 2013; Takahashi et al. 2017; Barth et al. 2020).

We investigated other impacts on the parental species based on the comparison with other *A. sandaracinos* and *A. chrysopterus* populations either occurring in sympatry (Solomon Islands) or allopatry (respectively Christmas Islands and Fiji; Figure 5B). Patterns of *FST* were highly similar among the three pairwise comparisons. Nucleotide diversity varied among populations, and the level was generally lower in *A. sandaracinos* compared to *A. chrysopterus*, consistent with what was previously highlighted using microsatellite markers (Gainsford et al. 2015). However, we found a considerable difference in the *dxy* distribution when comparing sympatric and allopatric populations. Surprisingly, *dxy* – the absolute genetic divergence – was lower between the allopatric populations (*A. sandaracinos* from Christmas Islands – *A. chrysopterus* from Fiji), and all comparisons involving one of the allopatric populations (Supp. Figure S4). The genetic divergence was thus lower when comparing more geographically distant parental populations. Like various diversity measures, *dxy* is highly impacted by background selection, mutation rates and recombination rates across the genome (Charlesworth et al. 1993). Since such processes are usually conserved across related species – even after the completion of lineage sorting (Dutoit et al. 2017) – they cannot explain the observed *dxy* patterns. Due to the clownfishes’ propensity to hybridise (Schmid et al. 2022; Marcionetti and Salamin 2023), we cannot exclude that past hybridisation events involving other clownfish species occurred in the allopatric *A. chrysopterus* and *A. sandaracinos* parental populations. Indeed, the limited admixture exhibited by the *A. chrysopterus* population from Fiji was unexpected (Figure 1) and intriguingly similar to the ancestry patterns previously observed in the skunk clownfish complex – whose evolutionary history was marked by ancestral genetic exchange (Schmid et al. 2022; Marcionetti and Salamin 2023). A sister species to *A. sandaracinos* – *A. perideraion –* occurs in both allopatric locations and has a history of past hybridisation with other clownfish species, notably with *A. sandaracinos* (Schmid et al. 2022, Marcionetti and Salamin 2023). We thus carefully suggest that the reduced divergence between the allopatric parental populations might be a consequence of past genetic exchange with the clownfish *A. perideraion*, and is not linked to the dynamics of the Kimbe Bay hybrid zone. The subsequent split in the admixture plots of the Fiji population from the other *A. chrysopterus* populations (at *K* = 3) followed by the separation of the Christmas Islands *A. sandaracinos* from other conspecific populations (at *K* = 4; Supp. Figure S1) further support that distinct evolutionary processes might have shaped the genome of the two parental allopatric populations compared to the sympatric ones.

### Future directions

The recurrent backcrossing of the F1-hybrids with *A. sandaracinos* combined with the scarcity of second-generation hybrids and the genomic signature of past genetic exchange among the parental species suggest that the hybridisation events occurring in the Kimbe Bay hybrid zone might lead to adaptive introgression. The resulting stabilised hybrid genome would thus consist of a few potentially *A. chrysopterus* adaptive loci introgressing into the *A. sandaracinos* genomic background. However, we expect distinct outcomes at other locations of the *A. chrysopterus-A. sandaracinos* hybrid zone – which ranges from the Solomon Islands to the Halmahera Island. Indeed, backcrosses were detected in Kimbe Bay but not in other locations (i.e. Solomon Islands and Kavieng), which consisted mainly of equally admixed individuals (Gainsford et al. 2020). Dissimilar hybridisation outcomes also occurred in the rainbow trout hybrid zone, where some locations only consisted of advanced backcross hybrids, while others were constituted of a mix between F1-hybrids and backcrosses (Mandeville et al. 2019). Geographical heterogeneity in hybridisation outcomes was also highlighted in Catostomus suckers (Mandeville et al. 2015; 2017) or in the hybrid zone between the rainbow and the westslope trout (Young et al. 2016; Muhlfeld et al. 2017). This spatial variation of introgression patterns and hybrid frequency is a consequence of the variability among locations in multiple processes such as reproductive isolation (Teeter et al. 2010; Gompert et al. 2014; Mandeville et al. 2017), context-specific selection or drift (Ottenburghs et al. 2017).

We thus expect a high variability in the hybridisation outcomes within the *A. chrysopterus-A. sandaracinos* hybrid zone, which we might be able to highlight by proceeding to the genomic analysis of more locations. Such genomic data would provide the substrate to answer some outstanding questions such as: to what extent different parental combinations can be achieved? What is the relative importance of locally adapted variances compared to the constraints imposed by the recombination rate? Such knowledge should lead to a deeper understanding of the mechanisms linked with clownfish speciation and reproductive isolation.

## MATERIAL AND METHODS

### Sampling, library preparation and sequencing

Between 2011 and 2014, we collected tissue samples consisting of 1 cm-long dorsal fin clips and took pictures from 23 *Amphiprion chrysopterus*, 24 *Amphiprion leucokranos* and 20 *Amphiprion sandaracinos*. All *A. leucokranos* (hybrid species) came from the hybrid zone in Kimbe Bay (Papua New Guinea). For the parental species, we collected *A. chrysopterus* and *A. sandaracinos* in the hybrid zone and in allopatric populations (Figure 1, Supp. Table S1).

We extracted the genomic DNA of each fin clip following the DNeasy Blood and Tissue kit standard procedure and performed the final elution twice in 100 µl of AE buffer (QIAGEN, Hombrechtikon, Switzerland). We quantified the extracted DNA using Qubit® 2.0 Fluorometer (Thermo Fisher Scientific, Waltham, Massachusetts, USA) and evaluated the integrity by electrophoresis. We followed the TruSeq Nano DNA library prep standard protocol to generate libraries with a 350 base pair insert size for whole-genome paired-end sequencing (Illumina, San Diego, California, USA). We validated the fragment length distribution of the libraries with a fragment analyser (Agilent Technologies, Santa Clara, California, USA). We pooled all samples together, and the Genomic Technologies Facility of the University of Lausanne performed the sequencing on ten lanes of Illumina 4000 HiSeq to reach an approximate 10X coverage for each sample.

### Sequenced data processing, mapping and genotyping

We trimmed the generated reads for adapter sequences using Cutadapt V2.3(Martin 2011). We filtered reads shorter than 40 bp and with a Phred quality score below with Sickle V1.33 (Joshi and Fass 2011). We assessed read quality before and after processing with FastQC V0.11.7 (Andrews 2010). We mapped the processed reads to *Amphiprion percula* reference genome (GenBank Assembly ID GCA_003047355.1; Lehmann et al. 2018) using BWA-MEM V0.7.17 (Li et al. 2009) and subsequently sorted, indexed and filtered them according to mapping quality (>30) using SAMtools V1.8 (Li et al. 2009). Then, we assigned all the reads to read-groups using Picard Tools V2.20.7 (http://broadinstitute.github.io/picard/), and we merged overlapping reads using ATLAS V0.9 (Link et al. 2017). We validated the mapping output using various statistics generated with BamTools V2.4.1 (Barnett et al. 2011). After mapping, we computed genotype likelihood and estimated the major and minor alleles using the Maximum Likelihood Estimation method using ATLAS V0.9 (Link et al. 2017). We filtered the resulting VCF file with VCFtools V0.1.14 (Danecek et al. 2011) based on mean minimum site depth (--minDP 2), mean maximum site depth (--maxDP 50), minor allele frequency (--maf 0.06), minimum quality site (--QUAL 40) and included only sites informative for all individuals.

### Principal component analysis and admixture

To explore the relationship between the two parental species and the hybrid, we first performed a principal component analysis (PCA) including all samples and based on the covariance matrix between individuals with PCAngsd V0.981 (Meisner and Albrechtsen 2018). Then, we estimated individual admixture proportions using NGSadmix V32 (Skotte et al. 2013) with *K* values ranging between two and seven. To determine the best number of *K* ancestral populations, we ran each NGSadmix analysis five times and calculated the probability of each *K* based on the likelihood values of the five runs following Evanno et al. (2005) procedure. To further investigate the relationship between the hybrids, we performed a second PCA analysis, including only the *A. leucokranos* (hybrid species) individuals.

### Hybrid index

To ensure consistency between methods to determine the ancestry proportion of hybrids, we calculated the proportion of alleles copies coming from each parental reference species to estimate the hybrid index (HI) using the Bayesian algorithm implemented in the R package GGHYBRID V1.0.0 (Buerkle 2005; Ribailey 2022). We considered as parental species *A. chrysopterus* and *A. sandaracinos* coming from the different sampled populations and based the analysis on a subset of SNPs. We pruned the SNP dataset for markers in linkage disequilibrium (LD) using a *r^2^ = 0.2* threshold and subsequently selected the 10% most differentiated SNPs based on pairwise *FST* calculation among the two parental species. We implemented further filtering within the GGHYBRID framework and considered only loci with parental allele frequency difference above 0.2 and removed loci with a parental species overlap in the Bayesian posterior 95% credible intervals of allele frequency. We computed the HI estimations using 3,000 Markov Chain Monte Carlo (MCMC) iterations after a 1,000 MCMC burn-in period, as suggested in the GGHYBRID R package.

### Mitochondrial genome reconstruction and phylogeny

To explore the genetic relationship between individuals and determine the parental inheritance of mitochondrial DNA in hybrids, we first reconstructed the mitochondrial genome of the 67 individuals using the baiting and iterative mapping approach implemented in MITObim V1.6 (Hahn et al. 2013). We applied two different reconstruction methods and checked for consistency between the outputs. We either (1) used the complete mitochondrial genome of *A. percula* (GenBank accession: KJ174497.1; Tao et al. 2016) as reference or (2) used *A. chrysopterus* and *A. sandaracinos* COI barcode sequences (GenBank accessions: FJ582751.1 and FJ582814; Steinke et al. 2009) as starting seed. We circularised the 67 resulting genomes with MARS (Ayad and Pissis 2017) and aligned them using MAFFT V7.475 (Katoh and Standley 2013). We then used the multiple sequences alignment as input to build a maximum-likelihood tree with IQ-tree V2.0.6 (Minh et al. 2020) and selected the substitution model (TPM2+F+R2) that minimises the Bayesian information criterion (BIC) with ModelFinder (Kalyaanamoorthy et al. 2017). We assessed branch support by implementing the single-branch test (SH-like approximate likelihood ratio; Guindon et al. 2010) and the ultrafast bootstrap approximation (UFBoot; Hoang et al. 2018) with 1,000 replicates and eventually visualise the resulting phylogeny with iTOL V6 (Letunic and Bork 2021).

### Local ancestry inference

We used the two-layer hidden Markov model implemented in ELAI (Guan 2014) to infer the local ancestry of the hybrid individuals. This approach estimates linkage disequilibrium (LD) between predefined parental groups to calculate the allele dosage scores at each SNP. The values range from 0 to 2 in a two-way admixture and indicate the site’s most likely ancestry proportion (0 or 2 indicates homozygous states, whereas a value of 1 corresponds to a heterozygous state). We ran ELAI on the complete SNPs dataset and considered the allopatric populations of *A. chrysopterus* (Fiji) and *A. sandaracinos* (Christmas Islands) as the two parental groups. We set the number of upper-layer clusters (-C) to 2 (representing *A. sandaracinos* and *A. chrysopterus*) and the lower-layer clusters (-c) to 10 (five times the upper-layer cluster, following ELAI recommendations). We applied six different admixture generation (-mg) parameters (1, 2, 3, 5, 10, 20) and performed ELAI analysis five times with 30 expectation maximization (EM) runs (-s) for each generation. We averaged the 30 independent runs and considered the sites with allele dosage between 0.8 and 1.2 as heterozygous, sites with allele dosage above 1.8 as homozygous for *A. chrysopterus* ancestry and sites below 0.2 as homozygous for *A. sandaracinos* ancestry, similar to what Seixas et al. (2018) suggested. We regarded values in-between those ranges as non-informative due to the high level of uncertainty.

### Hybrid colour patterns matrix and principal component analysis

The parental species differ in their phenotype in terms of white bands and background colour patterns: *A. chrysopterus* displays two white lateral bands and dark-orange background colouration, whereas *A. sandaracinos* has a single dorsal white stripe on a light-orange background (Figure 1C). The hybrids (*A. leucokranos* individuals) exhibit various intermediate traits between the two parental species. We built a colour pattern matrix consisting in nine binary traits (either absent or present in the hybrid): four traits describing the extent of the dorsal white band (S1-S4), four traits depicting the extent of the lateral white band (C1-C4) and one trait describing the asymmetry of pattern between the right and left side (A; Figure 2C). We thus characterised the 20 hybrid individuals with available pictures according to the presence or absence of those nine traits. Finally, we performed a principal component analysis (PCA) on the resulting matrix to summarise the colour information in a two-dimensional space and explore the relationship between the hybrids.

### Population genomic analyses in sliding windows

Based on the admixture analysis and the local ancestry estimation, we defined four groups of *A. leucokranos*: first-generation hybrids (F1), second-generation hybrids (F2), *sandaracinos*-backcrosses (BC) and *chrysopterus*-backcrosses (see Supp. Table S1). We only considered pure F1 (17 individuals) and *sandaracinos*-backcrosses (four individuals) for subsequent analyses. In addition, we removed the admixed *A. sandaracinos* (four individuals) from further analyses.

To investigate the genome-wide patterns of differentiation, divergence and diversity among the hybrids and the two parental populations, we estimated between-population differentiation (*FST*), between-population absolute divergence (*dxy*), as well as population nucleotide diversity (*π*) in non-overlapping 50kb windows (containing at least 100 SNPs) using popgenWindows.py (https://github.com/simonhmartin/genomics_general). We considered as populations the eight following groups: (1) F1 *A. leucokranos*, (2) BC *A. leucokranos*, (3-5) *A. sandaracinos* from Solomon Islands, Kimbe Bay and Christmas Islands and (6-8) *A. chrysopterus* from Solomon Islands, Kimbe Bay and Fiji (Supp. Table S1). Finally, we quantified the extent of genetic exchanges between the two parental lineages in the hybrid zone (Kimbe Bay) and calculated the *fdM* statistic (Malinsky et al. 2015) in non-overlapping 50kb windows (containing at least 100 SNPs) using popgenWindows.py. Like other *f-*statistics (also known as ABBA-BABA statistics), the *fdM* statistic relies on patterns of ancestral and derived alleles in a group of species/populations of interest (P1, P2 and P3) and an outgroup to discern hybridisation from incomplete lineage sorting. Compared to other similar statistics, *fdM* has the additional advantage of accommodating genetic exchanges between both P2 and P3 (positive value) and P1 and P3 (negative values). Since we were interested in genetic exchanges between parental species in Kimbe Bay, we considered the following topologies: P1 = *A. sandaracinos* (Christmas Islands), P2 = *A. sandaracinos* (Kimbe Bay), P3 = *A. chrysopterus* (Kimbe Bay) and outgroup = *A. chrysopterus* (Fiji).

We subsequently used those statistics to characterise the genomic landscapes of the F1-hybrids, the *sandaracinos*-BC hybrids and the parental lineages based on the following analyses.

#### Identification of potential barrier loci in parental populations

To identify potential barrier loci among the parental species *A. chrysopterus* and *A. sandaracinos*, we used the pairwise *FST* between the two species in sympatry (the Solomon Islands and Kimbe Bay) and in allopatry (*A. chrysopterus* from Fiji and *A. sandaracinos* from Christmas Islands). Regions of the genome with elevated *FST* can point to fixed or nearly fixed allele frequency differences between populations (i.e. barrier loci/genomic islands of differentiation). We thus extracted the 50kb genomic windows with *FST* values above the 99^th^ percentile of the whole-genome distribution and in common between three pairwise comparisons. We subsequently considered those windows as potential barrier loci.

#### Identification of regions of elevated divergence between F1-hybrids and one of the parental species

We used *FST* disparities between lineages to identify genomic regions of inflated divergence between the F1-hybrids and either or both parental species (also referred in results and discussion as “outlier windows”). Thus, for each 50kb-window across the genome, we computed the difference in *FST* between the F1-hybrids and both parental species from Kimbe Bay and subsequently selected the *FST*-difference values above the 99^th^ percentile (biased toward *A. chrysopterus*) and below the 1^st^ percentile (biased toward *A. sandaracinos*; Eq. 1)

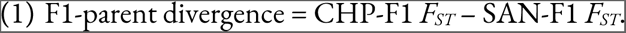

To detect if the number of highly divergent windows was biased toward some chromosomes, we generated a random distribution of highly divergent windows across the genome using 10,000 permutations. We then compared the observed mean value of divergent windows per chromosome to the expected mean value. We corrected the *p*-value for multiple testing using the Benjamini-Hochberg false-discovery rate method (Benjamini and Hochberg 1995). We performed a two-sample Wilcoxon test to evaluate differences in *FST*, *dxy* and *fdM* values between the background genomic windows and the highly divergent windows. Since the statistical significance indicated by the *p*-value is highly influenced by the sample size and can be misleading when many observations are compared (like genomic data), we computed the effect size using Cohen’s *d* statistic to assess the practical significance of the difference between the two groups. We also performed a chi-square test to compare the number of barrier loci between background and divergent genomic windows. Finally, we performed a one-sample Wilcoxon rank-sum test to test whether the mean divergence value across chromosomes was statistically different from 0.

#### Characterisation of introgressed A. sandaracinos tracts in BC A. leucokranos

Similar to the analyses we made for the divergent windows between F1-hybrids and the parental species, we used the population genomic statistics to characterise the introgressed *A. sandaracinos* genomic tracts identified with ELAI in the *A. leucokranos* backcrosses.

Here, we considered the windows with at least four out of the five *A. leucokranos* backcrosses displaying *A. sandaracinos* ancestry to be potential introgressed tracts. To test for a bias in some chromosomes compared to the rest of the genome, we generated a random distribution of introgressed windows across the genome using 10,000 permutations. We then compared the observed mean value of introgressed windows per chromosome to the expected mean value. We corrected the *p*-value for multiple testing using the Benjamini-Hochberg false-discovery rate method (Benjamini and Hochberg 1995). Then, we tested for differences in the *FST*, *dxy* and *fdM* values between the background and introgressed genomic windows using a two-sample Wilcoxon test and computed the effect size using Cohen’s *d* statistic. Finally, we performed a chi-square test to compare the number of barrier loci between background and introgressed genomic windows.

### Gene ontology

We extracted (1) the genes located within the *A. sandaracinos* introgressed tracts in backcross individuals as well as (2) the genes in the highly divergent windows between F1-hybrid and parental species to test for enrichment in specific gene ontology (GO) terms (done separately for both datasets). We used the 17,179 annotated genes inferred by Marcionetti *et al*. (2019), among which 14,002 were annotated with *biological process* GO terms. We retrieved gene annotations for all the above-mentioned genes of interest and performed GO enrichment analysis by comparing those annotations to the complete set of protein-coding annotated genes of *A. percula*. We used the R package topGO V2.44 (Alexa and Rahnenfuhrer 2020) to perform gene ontology enrichment analysis based on Fisher’s exact test with the *weight01* algorithm. We defined a minimum node size of 10 and considered as significant those GO terms with a raw *p*-value below 0.01, following the recommendations from the topGO manual. We reduced redundancy among GO terms with REVIGO (Supek et al. 2011) and clustered related GO terms according to their level of semantic similarity (>70% with SimRel similarity measure) using the *Danio rerio* database.

## DATA ACCESSIBILITY STATEMENT

Raw reads were deposited on NCBI (BioProject ID: PRJNA1078272) and will be made available upon acceptance of the manuscript. VCF files, scripts and images will be deposited on Dryad upon acceptance of the manuscript.

## AUTHOR CONTRIBUTIONS

DH, NS and SS planned the study. AG and GPJ collected the samples. SS performed lab work and analysed the genomic data. DH analysed the morphological data. SS drafted the manuscript. All authors contributed to the final version of the manuscript.

## ACKNOWLEDGEMENT

We thank the Mahonia Na Dari Research and Conservation Centre and Walindi Resort for supporting field work in Kimbe Bay. Collections were made in accordance with James Cook University animal ethics approval number A1633. We also would like to thank the Lausanne Genomic Technologies Facility for sequencing the samples and the DCSR infrastructure of the University of Lausanne for the computational resources. We are grateful to Anna Marcionetti for useful feedback on the manuscript and insightful discussions. Funding was from the University of Lausanne funds, Swiss National Science Foundation, Grant Number: 31003A-163428

## REFERENCES

Abbott RJ, Albach D, Ansell S, Arntzen JW, Baird SJE, Bierne N, Boughman J, Brelsford A, Buerkle CA, Buggs R et al. 2013. Hybridization and speciation. J. Evol. Biol. 26:229–246.

Abbott RJ. 2017. Plant speciation across environmental gradients and the occurrence and nature of hybrid zones. J. Syst. Evol. 55:238–258.

Alexa A, Rahnenfuhrer J. 2020. topGO: Enrichment analysis for Gene Ontology. R package version 2.43.0.

Allendorf FW, Leary RF, Spruell P, Wenburg JK. 2001. The problems with hybrids: setting conservation guidelines. Trends Ecol. Evol. 16:613–622.

Andrews S. 2010. FastQC: a quality control tool for high throughput sequence data. Available online at: http://www.bioinformatics.babraham.ac.uk/projects/fastqc/. Accessed July 6, 2020.

Ang TZ, Manica A. 2010. Unavoidable limits on group size in a body size-based linear hierarchy. Behav. Ecol. 21:819–825.

Arnegard ME, McGee MD, Matthews B, Marchinko KB, Conte GL, Kabir S, Bedford N, Bergek S, Chan YF, Jones FC, et al. 2014. Genetics of ecological divergence during speciation. Nature. 511:307–311.

Arnold ML. 1997. Natural Hybridization and Evolution. Oxford University Press: Oxford, New York.

Ayad LAK, Pissis SP. 2017. MARS: improving multiple circular sequence alignment using refined sequences. BMC Genom. 18.

Barnett DW, Garrison EK, Quinlan AR, Strömberg MP, Marth GT. 2011. BamTools: a C++ API and toolkit for analyzing and managing BAM files. Bioinformatics. 27:1691–1692.

Barth JMI, Gubili C, Matschiner M, Tørresen OK, Watanabe S, Egger B, Han Y-S, Feunteun E, Sommaruga R, Jehle R, et al. 2020. Stable species boundaries despite ten million years of hybridization in tropical eels. Nat. Commun. 11:1–13.

Barton NH, Hewitt GM. 1985. Analysis of hybrid zones. Annu. Rev. Ecol. Syst. 16:113–48.

Barton NH. 1986. The effects of linkage and density-dependent regulation on gene flow. Heredity. 57:415–426.

Barton NH, Bengtsson BO. 1986. The barrier to genetic exchange between hybridising populations. Heredity. 57:357–376.

Barton NH, Hewitt GM. 1989. Adaptation, speciation and hybrid zones. Nature. 341:497–503.

Barton NH. 2001. The role of hybridization in evolution. Mol Ecol. 10:551–568.

Barton NH, De Cara MAR. 2009. The evolution of strong reproductive isolation. Evolution. 63:1171–1190.

Barton NH. 2018. The consequences of an introgression event. Mol Ecol. 27:4973–4975.

Benjamini Y, Hochberg Y. 1995. Controlling the false discovery rate: a practical and powerful approach to multiple testing. J. R. Stat. Soc. B. 57:289–300.

Bolnick DI, Near TJ. 2005. Tempo of hybrid inviability in centrarchid fishes (Teleostei: Centrarchidae). Evolution. 59:1754–1767.

Brandvain Y, Kenney AM, Flagel L, Coop G, Sweigart AL. 2014. Speciation and Introgression between *Mimulus nasutus* and *Mimulus guttatus*. PLOS Genet. 10:e1004410.

Brelsford A, Irwin DE. 2009. Incipient speciation despite little assortative mating: the yellow-rumped warbler hybrid zone. Evolution. 63:3050–3060.

Buerkle CA. 2005. Maximum-likelihood estimation of a hybrid index based on molecular markers. Mol Ecol. Notes. 5:684–687.

Buerkle CA, Rieseberg LH. 2008. The rate of genome stabilization in homoploid hybrid species. Evolution. 62:266–275.

Burton RS. 1990. Hybrid breakdown in physiological response: a mechanistic approach. Evolution. 44:1806–1813.

Burton RS, Barreto FS. 2012. A disproportionate role for mtDNA in Dobzhansky–Muller incompatibilities? Mol Ecol. 21:4942–4957.

Burton RS, Ellison CK, Harrison JS. 2006. The sorry state of F2 hybrids: consequences of rapid mitochondrial DNA evolution in allopatric populations. Am. Nat. 168:S14–S24.

Buston PM. 2003. Size and growth modification in clownfish. Nature. 424:145–146.

Calfee E et al. 2021. Selective sorting of ancestral introgression in maize and teosinte along an elevational cline. PLOS Genet. 17:e1009810.

Carlon DB, Robertson DR, Barron RL, Choat JH, Anderson DJ, Schwartz SA, Sánchez-Ortiz CA. 2021. The origin of the parrotfish species *Scarus compressus* in the Tropical Eastern Pacific: region-wide hybridization between ancient species pairs. BMC Ecol. Evol. 21:7.

Charlesworth B, Morgan MT, Charlesworth D. 1993. The effect of deleterious mutations on neutral molecular variation. Genetics. 134:1289–1303.

Christie K, Strauss SY. 2018. Along the speciation continuum: quantifying intrinsic and extrinsic isolating barriers across five million years of evolutionary divergence in California jewelflowers. Evolution 72:1063–1079.

Danecek P, Auton A, Abecasis G, Albers CA, Banks E, DePristo MA, Handsaker RE, Lunter G, Marth GT, Sherry ST, et al. 2011. The variant call format and VCFtools. Bioinformatics 27:2156–2158.

Dobzhansky T. 1950. Genetics of natural populations. XIX. Origin of heterosis through natural selection in populations of Drosophila pseudoobscura. Genetics. 35:288–302.

Dutoit L, Vijay N, Mugal CF, Bossu CM, Burri R, Wolf J, Ellegren H. 2017. Covariation in levels of nucleotide diversity in homologous regions of the avian genome long after completion of lineage sorting. Proceedings of the Royal Society B: Biological Sciences 284:20162756.

Edmands S, Burton RS. 1999. Cytochrome C oxidase activity in interpopulation hybrids of a marine copepod: a test for nuclear-nuclear or nuclear-cytoplasmic coadaptation. Evolution. 53:1972–1978.

Elgvin TO, Trier CN, Tørresen OK, Hagen IJ, Lien S, Nederbragt AJ, Ravinet M, Jensen H, Sætre G-P. 2017. The genomic mosaicism of hybrid speciation. Sci. Adv. 3:e1602996.

Ellison CK, Burton RS. 2008. Genotype-dependent variation of mitochondrial transcriptional profiles in interpopulation hybrids. Proc. Natl. Acad. Sci. U.S.A. 105:15831–15836.

Endler JA. 1977. Geographic Variation, Speciation and Clines. Princeton University Press: Princeton, New Jersey.

Evanno G, Regnaut S, Goudet J. 2005. Detecting the number of clusters of individuals using the software structure: a simulation study. Mol Ecol. 14:2611–2620.

Fautin DG, Allen GR. 1997. Field Guide to Anemone Fishes and Their Host Sea Anemones. 2nd Revised Edition. Western Australian Museum.

Feder JL, Egan SP, Nosil P. 2012. The genomics of speciation-with-gene-flow. Trends Genet. 28:342–350.

Fontaine MC, Pease JB, Steele A, Waterhouse RM, Neafsey DE, Sharakhov IV, Jiang X, Hall AB, Catteruccia F, Kakani E, et al. 2015. Extensive introgression in a malaria vector species complex revealed by phylogenomics. Science 347:1258524.

Fricke HW. 1979. mating system, resource defence and sex change in the anemonefish *Amphiprion akallopisos*. Z. Tierpsychol. 50:313–326.

Gainsford A, van Herwerden L, Jones GP. 2015. Hierarchical behaviour, habitat use and species size differences shape evolutionary outcomes of hybridization in a coral reef fish. J. Evol. Biol. 28:205–222.

Gainsford A, Jones GP, Hobbs JA, Heindler FM, Herwerden L. 2020. Species integrity, introgression, and genetic variation across a coral reef fish hybrid zone. Ecol. Evol. 10:11998– 12014.

Gavrilets S. 1997. Hybrid zones with Dobzhansky-type epistatic selection. Evolution. 51:1027– 1035.

Gay L, Crochet P-A, Bell DA, Lenormand T. 2008. Comparing clines on molecular and phenotypic traits in hybrid zones: a window on tension zone models. Evolution. 62:2789– 2806.

Gompert Z, Lucas LK, Buerkle CA, Forister ML, Fordyce JA, Nice CC. 2014. Admixture and the organization of genetic diversity in a butterfly species complex revealed through common and rare genetic variants. Mol Ecol. 23:4555–4573.

Gompert Z, Buerkle CA. 2016. What, if anything, are hybrids: enduring truths and challenges associated with population structure and gene flow. Evol Appl. 9:909–923.

Grant V. 1981. Plant Speciation. Columbia University Press.

Guan Y. 2014. detecting structure of haplotypes and local ancestry. Genetics. 196:625–642.

Guindon S, Dufayard J-F, Lefort V, Anisimova M, Hordijk W, Gascuel O. 2010. New algorithms and methods to estimate maximum-likelihood phylogenies: assessing the performance of PhyML 3.0. Syst Biol. 59:307–321.

Hagberg L, Celemín E, Irisarri I, Hawlitschek O, Bella JL, Mott T, Pereira RJ. 2022. Extensive introgression at late stages of species formation: insights from grasshopper hybrid zones. Mol Ecol. 31:2384–2399.

Hahn C, Bachmann L, Chevreux B. 2013. Reconstructing mitochondrial genomes directly from genomic next-generation sequencing reads – a baiting and iterative mapping approach. Nucleic Acids Res. 41:e129–e129.

Haines ML, Luikart G, Amish SJ, Smith S, Latch EK. 2019. Evidence for adaptive introgression of exons across a hybrid swarm in deer. BMC Evol Biol. 19(1):199.

Hanemaaijer MJ, Collier TC, Chang A, Shott CC, Houston PD, Schmidt H, Main BJ, Cornel AJ, Lee Y, Lanzaro GC. 2018. The fate of genes that cross species boundaries after a major hybridization event in a natural mosquito population. Mol Ecol. 27:4978–4990.

Harrison RG. 1990. Hybrid zones: windows on evolutionary process. Oxford Surv Evol Biol. 7:69–128.

Harrison RG. 1993. Hybrid Zones and the Evolutionary Process. Oxford University Press.

Harrison RG, Larson EL. 2014. Hybridization, introgression, and the nature of species boundaries. J Hered. 105:795–809.

Hench K, Helmkampf M, McMillan WO, Puebla O. 2022. Rapid radiation in a highly diverse marine environment. Proc Natl Acad Sci USA. 119:e2020457119.

Hermansen JS, Sæther SA, Elgvin TO, Borge T, Hjelle E, Sætre G-P. 2011. Hybrid speciation in sparrows I: phenotypic intermediacy, genetic admixture and barriers to gene flow. Mol Ecol. 20:3812–3822.

Hermansen JS, Haas F, Trier CN, Bailey RI, Nederbragt AJ, Marzal A, Sætre G-P. 2014. Hybrid speciation through sorting of parental incompatibilities in Italian sparrows. Mol Ecol. 23:5831–5842.

Hewitt GM. 1988. Hybrid zones – natural laboratories for evolutionary studies. Trends Ecol. Evol. 3:158–167.

Hewitt GM. 2001. Speciation, hybrid zones and phylogeography – or seeing genes in space and time. Mol Ecol. 10:537–549.

Hill GE. 2016. Mitonuclear coevolution as the genesis of speciation and the mitochondrial DNA barcode gap. Ecology and Evolution. 6:5831–5842.

Hoang DT, Chernomor O, von Haeseler A, Minh BQ, Vinh LS. 2018. UFBoot2: Improving the Ultrafast Bootstrap Approximation. Mol. Biol. Evol. 35:518–522.

Hvala JA, Frayer ME, Payseur BA. 2018. Signatures of hybridization and speciation in genomic patterns of ancestry. Evolution. 72:1540–1552.

Janoušek V, Munclinger P, Wang L, Teeter KC, Tucker PK. 2015. Functional organization of the genome may shape the species boundary in the house mouse. Mol. Biol. Evol. 32:1208– 1220.

Jiggins CD, Mallet J. 2000. Bimodal hybrid zones and speciation. Trends Ecol. 15:250–255.

Jiggins CD, Naisbit RE, Coe RL, Mallet J. 2001. Reproductive isolation caused by colour pattern mimicry. Nature. 411:302–305.

Jiggins CD, Salazar C, Linares M, Mavarez J. 2008. Hybrid trait speciation and *Heliconius* butterflies. Phil. Trans. R. Soc. B. 363:3047–3054.

Joshi N, Fass J. 2011. Sickle: a sliding-window, adaptive, quality-based trimming tool for FastQ files. https://github.com/najoshi/sickle.

Kalyaanamoorthy S, Minh BQ, Wong TKF, von Haeseler A, Jermiin LS. 2017. ModelFinder: fast model selection for accurate phylogenetic estimates. Nat Methods. 14:587–589.

Katoh K, Standley DM. 2013. MAFFT multiple sequence alignment software version 7: improvements in performance and usability. Mol. Biol. Evol. 30:772–780.

Keller I et al. 2013. Population genomic signatures of divergent adaptation, gene flow and hybrid speciation in the rapid radiation of Lake Victoria cichlid fishes. Mol Ecol. 22:2848– 2863.

Keller L, Reeve HK. 1994. Partitioning of reproduction in animal societies. Trends Ecol Evol. 9:98–102.

Kirkpatrick M, Servedio MR. 1999. The reinforcement of mating preferences on an island. Genetics. 151:865–884.

Kondrashov AS. 2003. Accumulation of Dobzhansky-Muller incompatibilities within a spatially structured population. Evolution. 57:151–153.

Konkle BR, Philipp DP. 1992. Asymmetric hybridization between two species of sunfishes (*Lepomis: Centrarchidae*). Mol Ecol. 1:215–222.

Kovach RP, Hand BK, Hohenlohe PA, Cosart TF, Boyer MC, Neville HH, Muhlfeld CC, Amish SJ, Carim K, Narum SR, et al. 2016. Vive la résistance: genome-wide selection against introduced alleles in invasive hybrid zones. Proc. R. Soc. B 283:20161380.

Lai Z, Nakazato T, Salmaso M, Burke JM, Tang S, Knapp SJ, Rieseberg LH. 2005. extensive chromosomal repatterning and the evolution of sterility barriers in hybrid sunflower species. Genetics 171:291–303.

Lamb T, Avise JC. 1986. Directional introgression of mitochondrial DNA in a hybrid population of tree frogs: the influence of mating behavior. Proc. Natl. Acad. Sci. U.S.A. 83:2526–2530.

Lamichhaney S et al. 2017. Rapid hybrid speciation in Darwin’s finches. Science. eaao4593.

Lehmann R, Lightfoot DJ, Schunter C, Michell CT, Ohyanagi H, Mineta K, Foret S, Berumen M, Miller DJ, Aranda M, et al. 2018. Finding Nemo’s genes: a chromosome-scale reference assembly of the genome of the orange clownfish *Amphiprion percula*. Mol. Ecol. Resour. 19:570–585.

Lepais O et al. 2009. Species relative abundance and direction of introgression in oaks. Mol Ecol. 18:2228–2242.

Letunic I, Bork P. 2021. Interactive Tree Of Life (iTOL) v5: an online tool for phylogenetic tree display and annotation. Nucleic Acids Res. 49:W293–W296.

Li H, Handsaker B, Wysoker A, Fennell T, Ruan J, Homer N, Marth G, Abecasis G, Durbin R, 1000 Genome project data processing subgroup. 2009. the sequence alignment/map format and SAMtools. Bioinformatics 25:2078–2079.

Li Y, Tada F, Yamashiro T, Maki M. 2016. Long-term persisting hybrid swarm and geographic difference in hybridization pattern: genetic consequences of secondary contact between two *Vincetoxicum* species (Apocynaceae-Asclepiadoideae). BMC Evol Biol. 16:20.

Link V, Kousathanas A, Veeramah K, Sell C, Scheu A, Wegmann D. 2017. ATLAS: Analysis Tools for Low-depth and Ancient Samples. bioRxiv:105346. 10.1101/105346

Litsios G, Kostikova A, Salamin N. 2014. Host specialist clownfishes are environmental niche generalists. Proc. R. Soc. B. 281:20133220.

Maheshwari S, Barbash DA. 2011. The genetics of hybrid incompatibilities. Annu. Rev. Genet. 45:331–355.

Malinsky M, Challis RJ, Tyers AM, Schiffels S, Terai Y, Ngatunga BP, Miska EA, Durbin R, Genner MJ, Turner GF. 2015. Genomic islands of speciation separate cichlid ecomorphs in an East African crater lake. Science 350:1493–1498.

Mandeville EG, Parchman TL, McDonald DB, Buerkle CA. 2015. Highly variable reproductive isolation among pairs of *Catostomus* species. Mol Ecol. 24:1856–1872.

Mandeville EG, Parchman TL, Thompson KG, Compton RI, Gelwicks KR, Song SJ, Buerkle CA. 2017. Inconsistent reproductive isolation revealed by interactions between *Catostomus* fish species. Evol. Lett. 1:255–268.

Mandeville EG, Walters AW, Nordberg BJ, Higgins KH, Burckhardt JC, Wagner CE. 2019. Variable hybridization outcomes in trout are predicted by historical fish stocking and environmental context. Mol Ecol. 28:3738–3755.

Marcionetti A, Rossier V, Roux N, Salis P, Laudet V, Salamin N. 2019. insights into the genomics of clownfish adaptive radiation: genetic basis of the mutualism with sea anemones. Genome Biol Evol. 11:869–882.

Marcionetti A, Salamin N. 2023. Insights into the genomics of clownfish adaptive radiation: the genomic substrate of the diversification. Genome Biol Evol. 15:evad088.

Martin M. 2011. Cutadapt removes adapter sequences from high-throughput sequencing reads. EMBnet.journal. 17:10–12.

Martin SH, Davey JW, Salazar C, Jiggins CD. 2019. Recombination rate variation shapes barriers to introgression across butterfly genomes. PLoS Biol. 17:e2006288.

Mavárez J, Salazar CA, Bermingham E, Salcedo C, Jiggins CD, Linares M. 2006. Speciation by hybridization in *Heliconius* butterflies. Nature. 441:868.

McKenzie JL, Araújo HA, Smith JL, Schluter D, Devlin RH. 2021. Incomplete reproductive isolation and strong transcriptomic response to hybridization between sympatric sister species of salmon. Proc. R. Soc. B. 288:20203020.

Meisner J, Albrechtsen A. 2018. Inferring Population Structure and Admixture Proportions in Low-Depth NGS Data. Genetics. 210:719–731.

Melo MC, Salazar C, Jiggins CD, Linares M. 2009. assortative mating preferences among hybrids offers a route to hybrid speciation. Evolution. 63:1660–1665.

Minh BQ et al. 2020. IQ-TREE 2: new models and efficient methods for phylogenetic inference in the genomic era. Mol Biol Evol. 37:1530–1534.

Montanari SR, Hobbs J-PA, Pratchett MS, van Herwerden L. 2016. The importance of ecological and behavioural data in studies of hybridisation among marine fishes. Rev Fish Biol Fisheries. 26:181–198.

Muhlfeld CC, Kovach RP, Al-Chokhachy R, Amish SJ, Kershner JL, Leary RF, Lowe WH, Luikart G, Matson P, Schmetterling DA, et al. 2017. Legacy introductions and climatic variation explain spatiotemporal patterns of invasive hybridization in a native trout. Glob. Change Biol. 23:4663–4674.

Navascués M, Becheler A, Gay L, Ronfort J, Loridon K, Vitalis R. 2020. Power and limits of selection genome scans on temporal data from a selfing population. bioRxiv:2020.05.06.080895. 10.1101/2020.05.06.080895.

Nelson TC, Stathos AM, Vanderpool DD, Finseth FR, Yuan Y, Fishman L. 2021. Ancient and recent introgression shape the evolutionary history of pollinator adaptation and speciation in a model monkeyflower radiation (*Mimulus* section *Erythranthe*). PLoS Genet. 17:e1009095.

Nice CC, Gompert Z, Fordyce JA, Forister ML, Lucas LK, Buerkle CA. 2013. Hybrid speciation and independent evolution in lineages of alpine butterflies. Evolution. 67:1055– 1068.

Nolte AW, Gompert Z, Buerkle CA. 2009. Variable patterns of introgression in two sculpin hybrid zones suggest that genomic isolation differs among populations. Mol Ecol. 18:2615– 2627.

Orr HA. 1995. The population genetics of speciation: the evolution of hybrid incompatibilities. Genetics. 139:1805–1813.

Ostberg CO, Hauser L, Pritchard VL, Garza JC, Naish KA. 2013. Chromosome rearrangements, recombination suppression, and limited segregation distortion in hybrids between Yellowstone cutthroat trout (*Oncorhynchus clarkii bouvieri*) and rainbow trout (*O. mykiss*). BMC Genom. 14:570.

Ottenburghs J, Kraus RHS, van Hooft P, van Wieren SE, Ydenberg RC, Prins HHT. 2017. Avian introgression in the genomic era. Avian Res. 8:30.

Pardo-Diaz C, Salazar C, Baxter SW, Merot C, Figueiredo-Ready W, Joron M, McMillan WO, Jiggins CD. 2012. Adaptive introgression across species boundaries in *Heliconius* butterflies. PLOS Genet. 8:e1002752.

Payseur BA. 2010. Using differential introgression in hybrid zones to identify genomic regions involved in speciation. Mol. Ecol. Resourc. 10:806–820.

Poelstra JW, Vijay N, Bossu CM, Lantz H, Ryll B, Müller I, Baglione V, Unneberg P, Wikelski M, Grabherr MG, et al. 2014. The genomic landscape underlying phenotypic integrity in the face of gene flow in crows. Science. 344:1410–1414.

Presgraves DC. 2008. Sex chromosomes and speciation in Drosophila. Trends Genet. 24:336– 343.

Pujolar JM, Jacobsen MW, Als TD, Frydenberg J, Magnussen E, Jónsson B, Jiang X, Cheng L, Bekkevold D, Maes GE, et al. 2014. Assessing patterns of hybridization between North Atlantic eels using diagnostic single-nucleotide polymorphisms. Heredity. 112:627–637.

Pulido-Santacruz P, Aleixo A, Weir JT. 2018. Morphologically cryptic Amazonian bird species pairs exhibit strong postzygotic reproductive isolation. Proc. R. Soc. B. 285:20172081.

Rafati N, Blanco-Aguiar JA, Rubin CJ, Sayyab S, Sabatino SJ, Afonso S, Feng C, Alves PC, Villafuerte R, Ferrand N, et al. 2018. A genomic map of clinal variation across the European rabbit hybrid zone. Mol Ecol. 27:1457–1478.

Rand DM, Harrison RG. 1989. Ecological genetics of a mosaic hybrid zone: mitochondrial, nuclear, and reproductive differentiation of crickets by soil type. Evolution. 43:432–449.

Ravinet M, Yoshida K, Shigenobu S, Toyoda A, Fujiyama A, Kitano J. 2018. The genomic landscape at a late stage of stickleback speciation: High genomic divergence interspersed by small localized regions of introgression. Payseur BA, editor. PLoS Genet. 14:e1007358.

Ribailey RI. 2022. ribailey/gghybrid: gghybrid R package for Bayesian hybrid index and genomic cline estimation.

Rieseberg LH, Sinervo B, Linder CR, Ungerer MC, Arias DM. 1996. Role of gene interactions in hybrid speciation: evidence from ancient and experimental hybrids. Science. 272:741–745.

Rieseberg LH. 2001. Chromosomal rearrangements and speciation. Trends Ecol. 16:351–358.

Rieseberg LH, Raymond O, Rosenthal DM, Lai Z, Livingstone K, Nakazato T, Durphy JL, Schwarzbach AE, Donovan LA, Lexer C. 2003. Major ecological transitions in wild sunflowers facilitated by hybridization. Science. 301:1211–1216.

Runemark A, Trier CN, Eroukhmanoff F, Hermansen JS, Matschiner M, Ravinet M, Elgvin TO, Sætre G-P. 2018a. Variation and constraints in hybrid genome formation. *Nat*. Ecol. Evol. 2:549–556.

Runemark A, Eroukhmanoff F, Nava-Bolaños A, Hermansen JS, Meier JI. 2018b. Hybridization, sex-specific genomic architecture and local adaptation. Phil. Trans. R. Soc. B. 373:20170419.

Runemark A, Vallejo-Marin M, Meier JI. 2019. Eukaryote hybrid genomes. PLoS Genet. 15:e1008404.

Sankararaman S, Mallick S, Dannemann M, Prüfer K, Kelso J, Pääbo S, Patterson N, Reich D. 2014. The genomic landscape of Neanderthal ancestry in present-day humans. Nature. 507:354.

Sankararaman S, Mallick S, Patterson N, Reich D. 2016. The combined landscape of denisovan and neanderthal ancestry in present-day humans. Curr. Biol. 26:1241–1247.

Schaefer J, Duvernell D, Campbell DC. 2016. Hybridization and introgression in two ecologically dissimilar *Fundulus* hybrid zones. Evolution. 70:1051–1063.

Schmid S, Micheli B, Cortesi F, Donati GFA, Salamin N. 2022. Extensive hybridisation throughout clownfishes evolutionary history. bioRxiv: 2022.07.08.499304. 10.1101/2022.07.08.499304.

Schumer M, Rosenthal GG, Andolfatto P. 2014a. How common is homoploid hybrid speciation? Evolution. 68:1553–1560.

Schumer M, Cui R, Powell DL, Dresner R, Rosenthal GG, Andolfatto P. 2014b. High-resolution mapping reveals hundreds of genetic incompatibilities in hybridizing fish species. McVean G, editor. eLife. 3:e02535.

Schumer M, Cui R, Powell DL, Rosenthal GG, Andolfatto P. 2016. Ancient hybridization and genomic stabilization in a swordtail fish. Mol Ecol. 25:2661–2679.

Schumer M, Xu C, Powell DL, Durvasula A, Skov L, Holland C, Blazier JC, Sankararaman S, Andolfatto P, Rosenthal GG, et al. 2018. Natural selection interacts with recombination to shape the evolution of hybrid genomes. Science 360:656–660.

Schwarzbach AE, Donovan LA, Rieseberg LH. 2001. Transgressive character expression in a hybrid sunflower species. Am. J. Bot. 88:270–277.

Seixas FA, Boursot P, Melo-Ferreira J. 2018. The genomic impact of historical hybridization with massive mitochondrial DNA introgression. Genome Biol. 19:91.

Selz OM, Thommen R, Maan ME, Seehausen O. 2014. Behavioural isolation may facilitate homoploid hybrid speciation in cichlid fish. *Journal of Evol*. Biol. 27:275–289.

Servedio MR. 2004. The what and why of research on reinforcement. PLoS Biol. 2:e420.

Shurtliff QR. 2013. Mammalian hybrid zones: a review. Mammal Rev. 43:1–21.

Šimková A, Civáňová K, Vetešník L. 2022. Heterosis versus breakdown in fish hybrids revealed by one-parental species-associated viral infection. Aquaculture. 546:737406.

Simon A, Bierne N, Welch JJ. 2018. Coadapted genomes and selection on hybrids: Fisher’s geometric model explains a variety of empirical patterns. Evol. Lett. 2:472–498.

Singhal S, Moritz C. 2012. Strong selection against hybrids maintains a narrow contact zone between morphologically cryptic lineages in a rainforest lizard. Evolution. 66:1474–1489.

Skotte L, Korneliussen TS, Albrechtsen A. 2013. Estimating individual admixture proportions from next-generation sequencing data. Genetics. 195:693–702.

Sobel JM, Stankowski S, Streisfeld MA. 2019. Variation in ecophysiological traits might contribute to ecogeographic isolation and divergence between parapatric ecotypes of *Mimulus aurantiacus*. *Journal of Evol*. Biol. 32:604–618.

Stankowski S, Sobel JM, Streisfeld MA. 2015. The geography of divergence with gene flow facilitates multitrait adaptation and the evolution of pollinator isolation in *Mimulus aurantiacus*. Evolution. 69:3054–3068.

Stebbins GL. 1959. The role of hybridization in evolution. Proc. Am. Philos. Soc. 103:231–251.

Steinke D, Zemlak TS, Hebert PDN. 2009. Barcoding Nemo: DNA-based identifications for the ornamental fish trade. PLoS ONE. 4:e6300.

Stelkens RB, Schmid C, Seehausen O. 2015. Hybrid breakdown in cichlid fish. PLoS ONE. 10:e0127207.

Supek F, Bošnjak M, Škunca N, Šmuc T. 2011. REVIGO summarizes and visualizes long lists of gene ontology terms. PLoS ONE. 6:e21800.

Svedin N, Wiley C, Veen T, Gustafsson L, Qvarnström A. 2008. Natural and sexual selection against hybrid flycatchers. Proc. R. Soc. B. 275:735–744.

Takahashi H, Toyoda A, Yamazaki T, Narita S, Mashiko T, Yamazaki Y. 2017. Asymmetric hybridization and introgression between sibling species of the pufferfish *Takifugu* that have undergone explosive speciation. Mar Biol. 164:90.

Tao Y, Li J-L, Liu M, Hu X-Y. 2016. Complete mitochondrial genome of the orange clownfish *Amphiprion percula* (Pisces: Perciformes, Pomacentridae). Mitochondrial DNA A. 27:324–325.

Taylor EB, Boughman JW, Groenenboom M, Sniatynski M, Schluter D, Gow JL. 2006. Speciation in reverse: morphological and genetic evidence of the collapse of a three-spined stickleback (*Gasterosteus aculeatus*) species pair. Mol Ecol. 15:343–355.

Teeter KC, Thibodeau LM, Gompert Z, Buerkle CA, Nachman MW, Tucker PK. 2010. The variable genomic architecture of isolation between hybridizing species of house mice. Evolution. 64:472–485.

Thompson KA. 2020. Experimental hybridization studies suggest that pleiotropic alleles commonly underlie adaptive divergence between natural populations. Am. Nat. 196:E16–E22.

Toews DPL, Taylor SA, Vallender R, Brelsford A, Butcher BG, Messer PW, Lovette IJ. 2016. Plumage genes and little else distinguish the genomes of hybridizing warblers. Curr. Biol. 26:2313–2318.

Trier CN, Hermansen JS, Sætre G-P, Bailey RI. 2014. Evidence for mito-nuclear and sex-linked reproductive barriers between the hybrid italian sparrow and its parent species. PLoS Genet. 10:e1004075.

Veller C, Edelman NB, Muralidhar P, Nowak MA. 2023. Recombination and selection against introgressed DNA. Evolution. 77:1131–1144.

Wagner DN, Curry RL, Chen N, Lovette IJ, Taylor SA. 2020. Genomic regions underlying metabolic and neuronal signaling pathways are temporally consistent in a moving avian hybrid zone. Evolution. 74:1498–1513.

Walsh J, Shriver WG, Olsen BJ, O’Brien KM, Kovach AI. 2015. Relationship of phenotypic variation and genetic admixture in the Saltmarsh–Nelson’s sparrow hybrid zone. The Auk. 132:704–716.

Wong MYL. 2011. Group size in animal societies: the potential role of social and ecological limitations in the group-living fish, *Paragobiodon xanthosomus*. Ethology. 117:638–644.

Xiong T, Mallet J. 2022. On the impermanence of species: the collapse of genetic incompatibilities in hybridizing populations. Evolution. 76:2498–2512.

Yang W., Feiner, N., Pinho, C., While, G. M., Kaliontzopoulou, A., Harris, D. J., Salvi, D., and Uller, T. (2021). Extensive introgression and mosaic genomes of Mediterranean endemic lizards. Nat. Comm. 12(1), Article 2762.

Young MK, Isaak DJ, McKelvey KS, Wilcox TM, Pilgrim KL, Carim KJ, Campbell MR, Corsi MP, Horan DL, Nagel DE, et al. 2016. Climate, demography, and zoogeography predict introgression thresholds in salmonid hybrid zones in Rocky Mountain streams. PLoS ONE. 11:e0163563.

Zhang B-L, Chen W, Wang Z, Pang W, Luo M-T, Wang S, Shao Y, He W-Q, Deng Y, Zhou L, et al. 2023. Comparative genomics reveals the hybrid origin of a macaque group. Sci.Adv. 9:eadd3580.

