## Supplemental figure S1 for "Ongoing hybridisation among clownfishes: the genomic architecture of the Kimbe Bay hybrid zone"

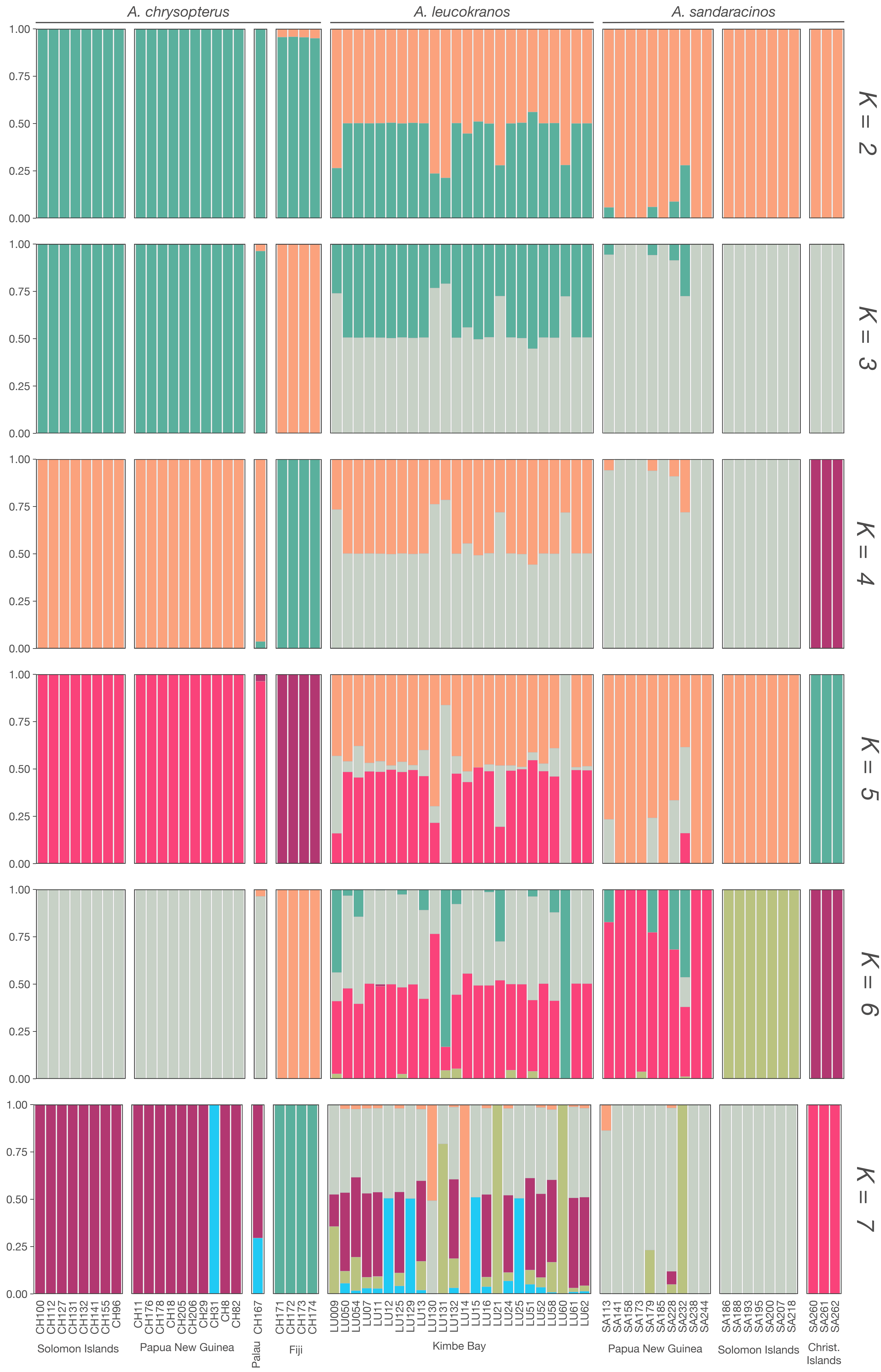


**Figure S1.** Admixture bar plots based on NGSadmix *q*-values for K = 2 to K = 7. Each bar represents the ancestry proportion of each individual for each cluster.
