## Supplemental figure S2 for "Ongoing hybridisation among clownfishes: the genomic architecture of the Kimbe Bay hybrid zone"

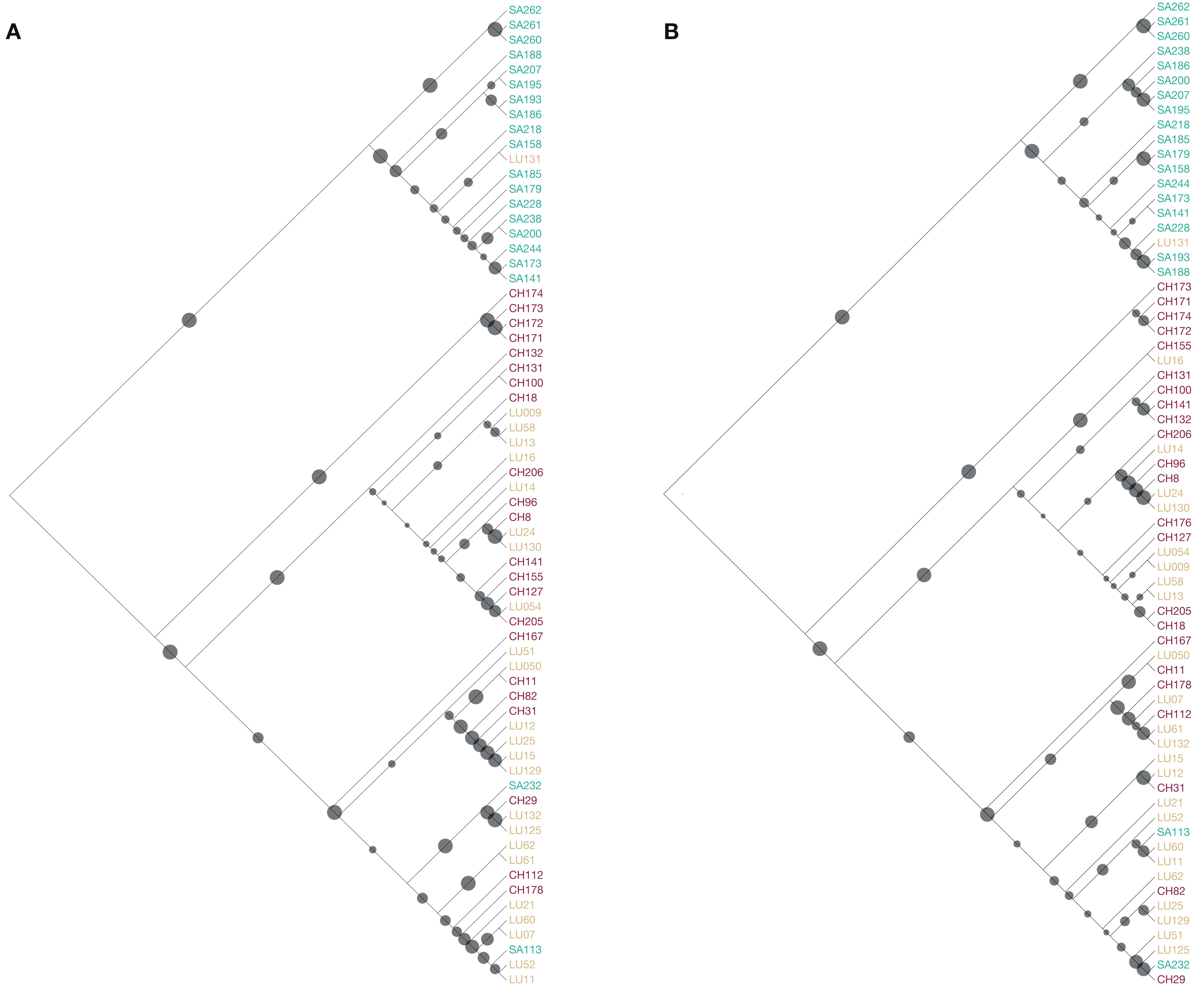


**Figure S2**. Mitochondrial midpoint-rooted phylogenetic reconstructed with IQ-tree V2.0.6. (A) The phylogeny is based on the 67 mitochondrial genomes reconstructed with MITObim using sandaracinos COI sequence as starting seed. The size of the dots on the branches corresponds to the bootstrap support (ranging from 2 to 100) based on 1,000 ultrafast bootstraps approximation. The colours correspond to the three species (dark red: *A. chrysopterus*; light brown: *A. leucokranos* (hybrid); turquoise: *A. sandaracinos*). (B) The phylogeny is based on the 67 mitochondrial genomes reconstructed with MITObim using *A. chrysopterus* COI sequence as starting seed. The size of the dots on the branches corresponds to the bootstrap support (ranging from 12 to 100) based on 1,000 ultrafast bootstraps approximation. The colours correspond to the three species (dark red: *A. chrysopterus*; light brown: *A. leucokranos* (hybrid); turquoise: *A. sandaracinos*).
