## Supplemental figure S3 for "Ongoing hybridisation among clownfishes: the genomic architecture of the Kimbe Bay hybrid zone"

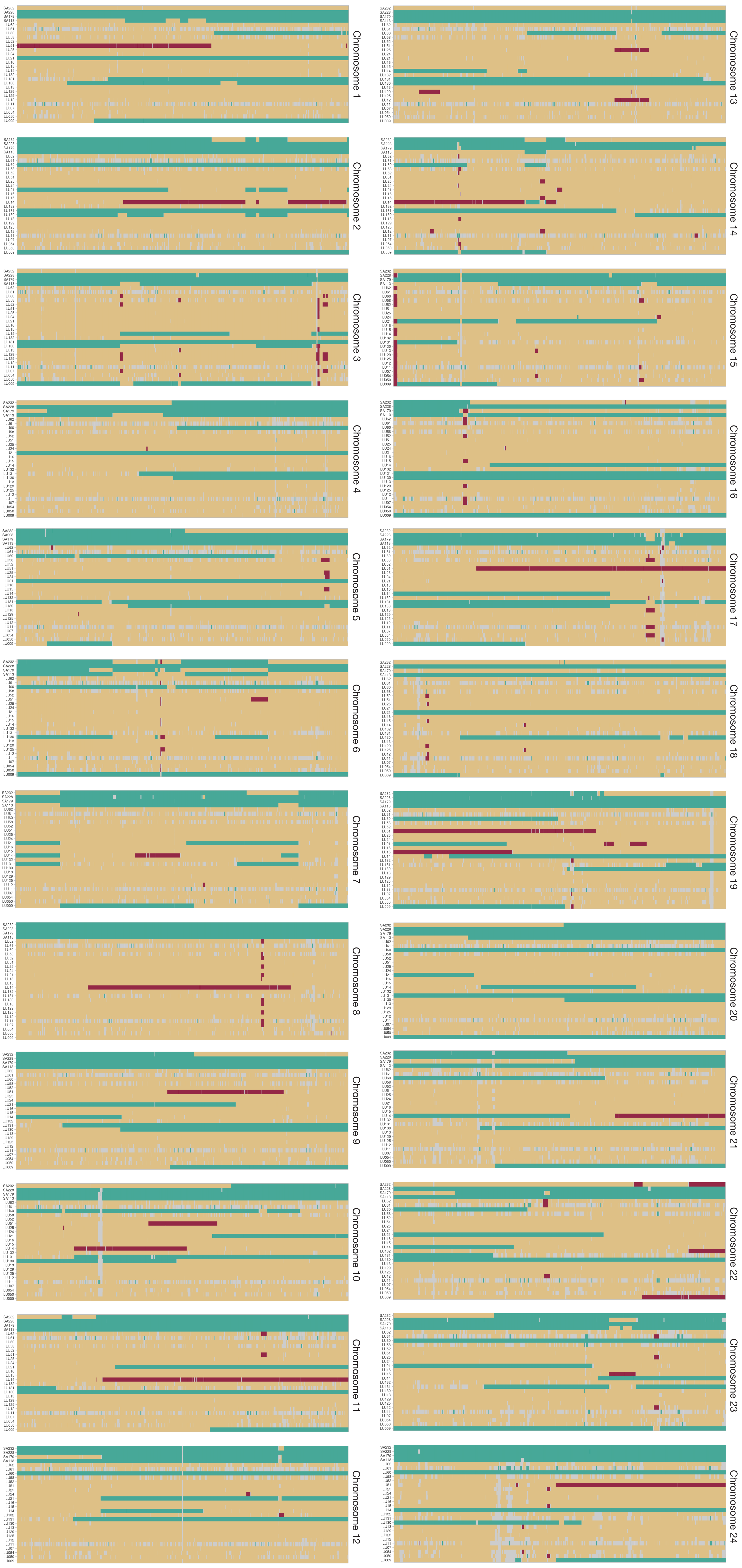


**Figure S3**. Ancestry estimated with ELAI (Guan 2014) across the genome for all *A. leucokranos* samples and the four admixed *A. sandaracinos*. Turquoise regions correspond to *A. sandaracinos* ancestry, dark red regions to *A. chrysopterus* ancestry, and the light brown colour highlights heterozygous regions. The grey colour indicates non-informative sites.
