## Supplemental figure S4 for "Ongoing hybridisation among clownfishes: the genomic architecture of the Kimbe Bay hybrid zone"

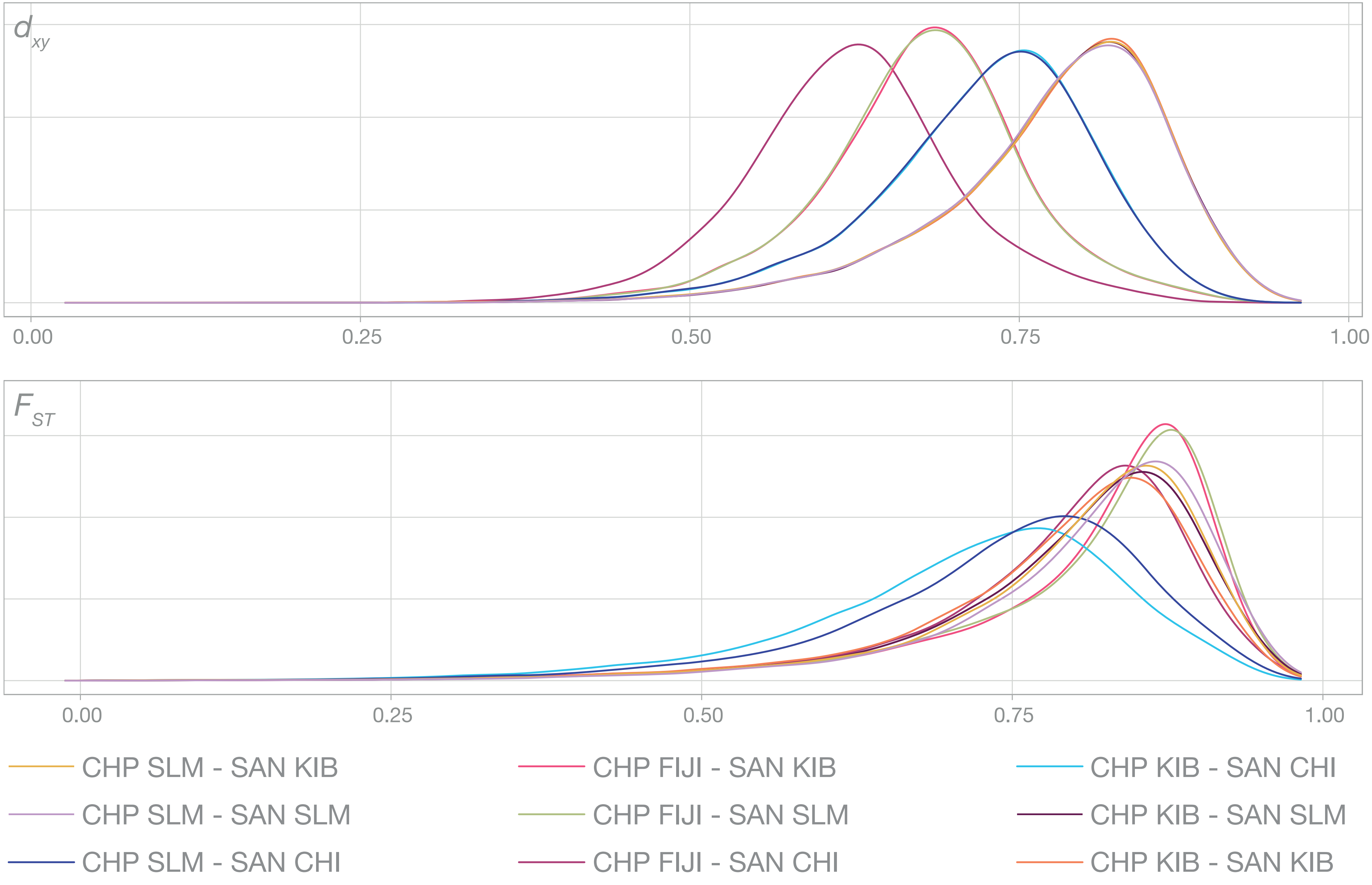


**Figure S4**. Density plot of $F_{ST}$ and $d_{xy}$, between all *A. sandaracinos* (SAN) and *A. chrysopterus* (CHP) populations sampled. CHI: Christmas Islands, FIJI: Fiji Islands, KIB: Kimbe Bay; SLM: Solomon Islands.
